# Evolution of genes involved in the unusual genitals of the bear macaque, *Macaca arctoides*

**DOI:** 10.1101/2020.05.18.102251

**Authors:** Laurie Stevison, Nick P Bailey, Zachary A Szpiech, Taylor E Novak, Don J Melnick, Ben J Evans, Jeffrey D Wall

**Affiliations:** Auburn University; Columbia University; McMaster University; University of California San Francisco

**Keywords:** reproductive isolation, hybridization, speciation, baculum

## Abstract

Genital divergence is thought to contribute to reproductive barriers by establishing a “lock-and-key” mechanism for reproductive compatibility. One such example, *Macaca arctoides*, the bear macaque, has compensatory changes in both male and female genital morphology as compared to close relatives. *Macaca arctoides* also has a complex evolutionary history, having extensive introgression between the *fascicularis* and *sinica* macaque species groups. Here, phylogenetic relationships were analyzed via whole genome sequences from five species, including *M. arctoides*, and two species each from the putative parental species groups. This analysis revealed ∼3x more genomic regions supported placement in the *sinica* species group as compared to the *fascicularis* species group. Additionally, introgression analysis of the *M. arctoides* genome revealed it is a mosaic of recent polymorphisms shared with both species groups. To examine the evolution of their unique genital morphology further, the prevalence of candidate genes involved in genital morphology were compared against genome-wide outliers in various population genetic metrics, while accounting for background variation in recombination rate. This analysis identified 66 outlier genes, including several genes that influence baculum morphology in mice, which were of interest since the bear macaque has the longest primate baculum. The mean of several metrics was statistically different in the candidate genes as compared to the rest of the genome, suggesting that genes involved in genital morphology have increased divergence and decreased diversity beyond expectations. These results highlight how extensive introgression may have contributed to reproductive isolation and shaped the unique genital morphology in the bear macaque.

## Introduction

In species with internal fertilization, genital morphology may have complex and rapid evolution. This has been extensively demonstrated in males (Eberhard, 1993; Klaczko *et al.*, 2015; Langerhans *et al.*, 2016) and females (Greenway *et al.*, 2019; Simmons & Fitzpatrick, 2019; Sloan & Simmons, 2019). There have been multiple proposed explanations for this including: (1) species isolation via a “lock-and-key” mechanism, (2) sexual conflict, and (3) cryptic female choice (Eberhard, 1985; Sloan & Simmons, 2019). The first explanation suggests that male and female genitalia uniquely coevolve to prevent heterospecific mating. It has historically been dismissed as a general explanation for genital evolution but acknowledged to play a role in particular cases (Eberhard, 1985, 2010). One reason for this dismissal is a presumed lack of variability in female genitalia between species, which suggests the “lock” (female genitalia) is rarely a unique match for the “key” (male genitalia) (Eberhard, 1985; Sloan & Simmons, 2019). However, studies in beetles (Sota & Kubota, 1998), flies (Kamimura & Mitsumoto, 2012), and damselflies (Barnard *et al.*, 2017) have demonstrated reproductive incompatibility between closely related species caused by mechanical mismatch in heterospecific crosses, suggestive of a “lock-and-key” mechanism (Sloan and Simmons, 2019). Sexual conflict may entail either conflict between males trying to obtain mates or conflict between males and females in levels of investment in offspring (Eberhard, 1985). Cryptic female choice happens when male reproductive output may be affected by the female after copulation (Eberhard, 1985, 1996). These processes both involve competition that may also result in species-specific genitalia. For example, a prediction of cryptic female choice is that female genitalia may evolve more rapidly than male genitalia. This can result in compensatory coevolution of male genitalia that causes reproductive isolation between species (Langerhans *et al.*, 2016; Greenway *et al.*, 2019; Sloan & Simmons, 2019). In support of this it has been found that male and female genitalia coevolve in internal fertilizing poecilid fishes (Greenway *et al.*, 2019) and in dung beetles female genitalia diverge more quickly than male genitalia (Simmons & Fitzpatrick, 2019). Additionally, it has been found that females in populations of dung flies exhibiting sexual conflict have greater preference for conspecific males than females in monogamous populations (Martin & Hosken, 2003).

The bear macaque, *Macaca arctoides*, is unique amongst macaques (and primates generally) for its genital morphology, with distinctive genital characteristics in both sexes (Figure 1). In males, it has the longest baculum of all primates (4-6 cm; Fooden, 1990), even when corrected for body weight (Dixson, 1987; Fooden, 1990). In addition to its elongation, the urethral opening is located on the ventral side of the penis, altering the overall morphology relative to other macaque species (Fooden, 1990). In females, *M. arctoides* female genitalia appear uniquely coadapted to their male counterparts in both length and morphology (Fooden, 1967). Given these compensatory changes, it has been hypothesized that genital evolution in this species followed a “lock-and-key” mechanism, which is likely to be rare amongst primates (Dixson, 1998).

**Figure 1.**
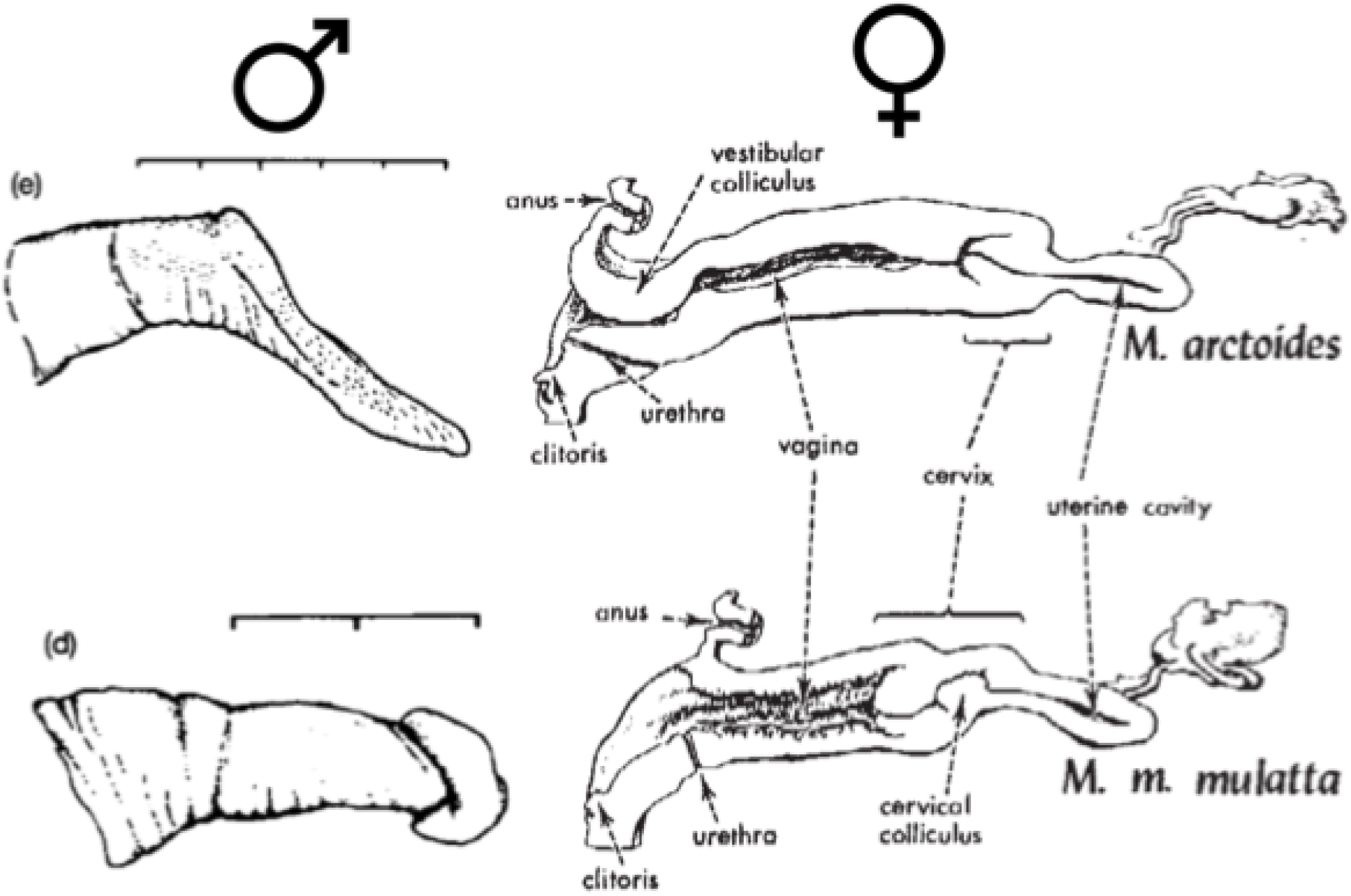
Diverged genital morphology in male and female bear macaques. *M. arctoides* has divergent male and female genital morphology (top), relative to M. mulatta (bottom) representing mechanical isolation between these species. M. mulatta morphology is representative of other species in macaques. Male external genitalia for each species are shown on the left (from Fig 3 of Dixson, 1987) and female reproductive tracts are shown on the right (from Fig 4 of Fooden, 1967). Reproduced with permission. Divergent traits for male genitalia include size and shape. Divergent traits for female reproductive tracts include depth of the uterine cavity, width of the vagina, opening size, and urethra location. Scales on the left are in 1 cm divisions.

Beyond genital morphology, the mating behavior of the bear macaque has been highlighted in the literature as being atypical amongst macaques. In most other macaques, males and females mount multiply, but perhaps due to the altered male genital morphology, bear macaques engage in a single-mount ejaculatory mating strategy with a prolonged sitting where the males deposit an ejaculatory plug in females (Fooden, 1990). However, most studies have excluded species from the *sinica* species group, except *M. arctoides* (Dixson, 1987; Dixson & Anderson, 2004). In studies looking at *M. radiata*, another member of the *sinica* species group, they also favor a single mount mating style differing from *M. fascicularis* and *M. mulatta* (Shively *et al.*, 1982). Similarly, to *M. arctoides*, *M. radiata* also exhibits a post-ejaculatory sit. Since both species have larger bacula than species in the *fascicularis* group, the hypothesis that increased baculum aide in prolonged intromission is supported. Therefore, the novelty of mating behavior in the bear macaque may be based on studies that did not include the behavior of representatives from its closest relatives.

Additional distinguishing characteristics not related to genital morphology include whitish pelage in newborns, and a bald forehead and cheeks (the latter evident in Figure 2A) (Fooden, 1990). Further, the evolutionary history of this species has been noted as extremely complex with early molecular work revealing phylogenetic incongruence between mitochondrial and Y-chromosomal genealogies for *M. arctoides* (Tosi *et al.*, 2000, 2003). Mitochondrial markers place *M. arctoides* as sister to the *fascicularis* species group (includes *M. cyclopis*, *M. fascicularis*, *M. fuscata*, and *M. mulatta*), whereas autosomal markers place it as sister to the *sinica* species group (includes M. assamensis, *M. radiata*, *M. sinica*, and M. thibetana) (Li *et al.*, 2009b; Jiang *et al.*, 2016; Li *et al.*, 2018; Fan *et al.*, 2018). Similarly, both morphological and Y-chromosomal trees support placement in the *sinica* group (Tosi *et al.*, 2000; Jiang *et al.*, 2016; Fan *et al.*, 2018) (see Table S1 in Fan *et al.*, 2018 for review). This phylogenetic incongruence was hypothesized to be consistent with ancient hybridization between the ancestor of the *fascicularis* species group and a *sinica* species group member (an ancestor of either *M. assamensis* or *M. thibetana*) contributing to hybrid speciation of *M. arctoides* (Tosi *et al.*, 2003).

**Figure 2.**
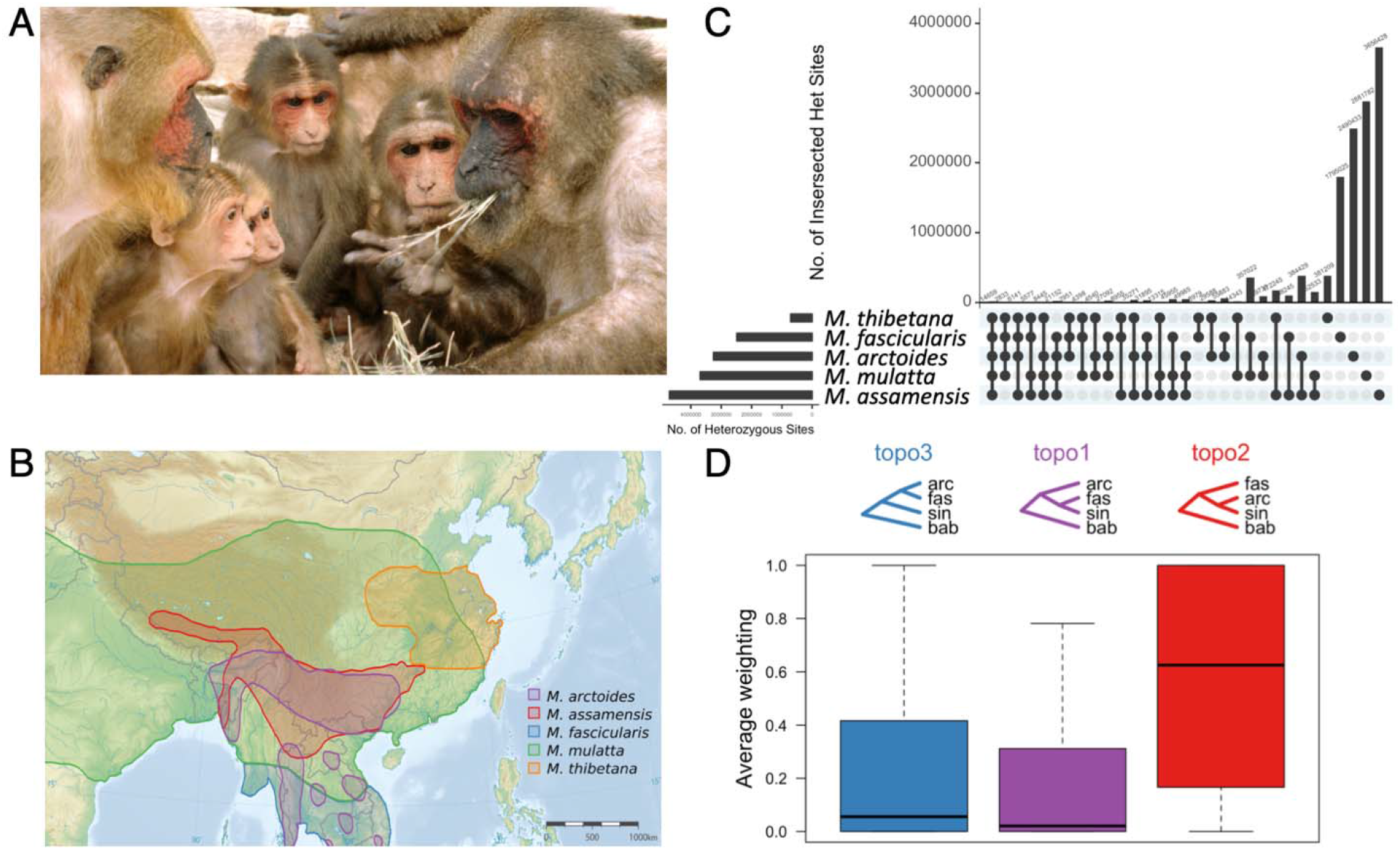
Summary of focal species, *Macaca arctoides*, its evolutionary relationships, and the data used. (a) *M. arctoides* has unique genital morphology and is a putative homoploid hybrid species. Image “Stumptail Monkeys” from (Waal, 2004) by Frans de Waal licensed under CC. (B) Geographic range of *M. arctoides* and other focal species in this study. *M. arctoides* has present day geographic overlap with members of both species groups (redrawn from (Fooden, 1980)). Background image “Topographic map of East Asia” by Ksiom is licensed under CC BY-SA 3.0. This study included WGS from *M. arctoides* and representative members of each species group (Table 1). Baboon was used as an outgroup species. (C) Intersection of heterozygous sites between the five species. The output of vcf-compare from vcftools (Danecek *et al.*, 2011) was input into the UpSetR package (Conway & Gehlenborg, 2019) to plot the intersection of sites. Values in the left plot match count of heterozygous sites per sample in Table 1. (D) Topology weight distribution of three rooted topologies of species relationships in sliding windows along the genome. bab - P. anubis.

**Table 1.** Sample information. A list of samples used in this study with a summary of genome information. Two samples were sequenced as part of this study (in bold) and the remainder were downloaded from NCBI.

| ID | species | Species Group | Coverage | No. sites VQSR Passed | No. sites heterozygous | NCBI Accession | SAMN_ID | Sex |
| --- | --- | --- | --- | --- | --- | --- | --- | --- |
| **Malaya** | ***M. arctoides*** | **arctoides** | **20** | **6,849,957** | **3,252,204** | **SRS6488501** | **14521207** | **F** |
| SM1 | *M. arctoides* | arctoides | 32 | 6,848,826 | 3,251,892 | SRR2981139 | 04316319 | F |
| SM2 | *M. arctoides* | arctoides | 19 | 6,833,185 | 3,246,931 | SRR2981140 | 04316320 | F |
| BGI-CE-4 | *M. fascicularis* | fascicularis | 43 | 4,547,767 | 2,486,762 | SRS114988 | 00116341 | F |
| CR-5 | *M. mulatta* | fascicularis | 44 | 4,561,581 | 3,693,024 | SRS115022 | 00113489 | F |
| BGI | *M. mulatta* | fascicularis | 10 | 4,531,187 | 3,678,452 | SRS212016 | 00627571 | M |
| BGI-96346 | *M. mulatta* | fascicularis | 4 | 4,334,465 | 3,533,580 | SRS114988 | 00113455 | M |
| **A20** | ***M. assamensis*** | **sinica** | **13** | **7,567,951** | **4,680,934** | **SRS6491954** | **14521608** | **F** |
| XH1 | *M. assamensis* | sinica | 49 | 7,669,854 | 4,702,055 | SRR2981114 | 04316321 | M |
| Tibetan-macaque-NO.3 | *M. thibetana* | sinica | 27 | 4,844,002 | 713,449 | SRR1024051 | 02390221 | F |

In contrast to allopolyploid speciation, homoploid hybrid speciation (HHS) conserves chromosome number and is postulated to occur with similar frequencies in plants and animals (Mallet, 2007; Mavárez & Linares, 2008). Though expected to be a rare speciation mechanism relative to bifurcating speciation, diverse taxa such as birds (*Passer italiae*; (Elgvin *et al.*, 2017)), sunflowers (Helianthus anomalus; (Rieseberg, 2003)) and yeast (*Saccharomyces paradoxus*; (Leducq *et al.*, 2016)) are thought to have evolved via HHS. Recent analysis of macaque genomes support introgression between *M. arctoides* and members of the *fascicularis* and *sinica* species groups (Fan *et al.*, 2018), both of whom it overlaps with geographically (Figure 2B). These species groups are estimated to have split ∼3 million years ago. Recent mitochondrial phylogenetics have shown that *M. arctoides* is more closely related to *M. mulatta* as compared to *M. fascicularis* (Fan *et al.*, 2018; Roos *et al.*, 2019), indicating introgression may have occurred after the split of these two species, which is estimated to be 1.68 mya (Stevison & Kohn, 2009; Jiang *et al.*, 2016; Fan *et al.*, 2018). This timing overlaps glacial periods of the Pleistocene which is associated with reduced expanse of Southeast Asian forests into multiple refugia. Ecological isolation is a common feature to other scenarios of HHS (Gompert *et al.*, 2006). In the case of *M. arctoides*, isolation is hypothesized to have contributed to the extensive interspecific hybridization forcing interbreeding of species that otherwise might have had alternative options for mating (Tosi *et al.*, 2003). Available evidence suggests that this interbreeding subsided ∼1.5 mya (Fooden, 1990; Tosi *et al.*, 2003). This hypothesized hybrid origin is diagrammed in Figure S1A, with a range of possible split times indicated by the thick purple line. Regardless of whether the bear macaque is indeed a hybrid species, the evidence for historical hybridization suggests that the dramatic, derived genital morphology may have been influenced by, or evolved despite extensive genetic exchange with other macaque species (Tosi *et al.*, 2000, 2003).

Here, we investigated the potential role of hybridization in shaping the unusual genital morphology of *M. arctoides*. We hypothesized that if the unique genital morphology contributes to reproductive isolation, then genes involved in genital morphology would show extreme population genetic signatures for *M. arctoides*. To test this hypothesis, we compiled a list of 2,284 candidate genes associated with genital morphology in other organisms, with a specific focus on mammals. We used an existing recombination rate map for *M. mulatta* to identify outliers in several population genetic metrics, such as introgression, nucleotide diversity, and divergence. Phylogenomic and population genetic approaches were used on several extant macaque species (*M. arctoides*; the *sinica* group species *M. thibetana* and *M. assamensis*; and the *fascicularis* group species *M. fascicularis* and *M. mulatta*). We found 66 of the candidate genes were significant genome outliers in at least one of seven population genetic metrics. Further, the mean across 4/7 population genetic metrics was significantly different in the candidate genes as compared to all other genes in the genome. Interestingly, three dN/dS outliers were associated with putative baculum morphology in mice. Overall, our results suggest that genital evolution in the bear macaque has been extensive and involved a large number of genes associated with various phenotypic changes in both males and females.

## Methods

### Samples and Genome Sequencing

One female each from *M. arctoides* (Malaya) and *M. assamensis* (A20) were sequenced. The *M. arctoides* sample was from the Malaka Zoo in Malaysia (previously described in Tosi *et al.*, 2000). Due to their now vulnerable conservation status, and restricted international trade, availability of these species is severely restricted, severely limiting sample sizes (see Discussion). These two samples were multiplexed and sequenced on one lane of sequencing for initial quality assessment. They were then subsequently run on one additional lane of sequencing each for a final 1.5 lanes per sample. Sequencing was done at UCSF Medical Center Sequencing Facility on an Illumina HiSeq 2000 machine. Each run was assigned as different read groups to be treated as independent in the variant calling workflow. The raw read data have been deposited in the NCBI sequence read archive and are available via the accession PRJNA622565 (see Table 1).

### Publicly available sequences

Raw FASTQ files were downloaded from 8 public genome samples for this project (Table 1). Genome samples were selected to be representative of the taxa relevant for *M. arctoides* evolution. For the *fascicularis* group, many more public sequences were available than used here, however we selected two sequencing projects and included all samples from both projects, including those with lower coverage (<10X, see Table 1). This sample size was chosen to allow for equal representation across the various species groups. Additional sample details can be found in the corresponding publications (Yan *et al.*, 2011; Fan *et al.*, 2014, 2018). These were downloaded from NCBI SRA using sratoolkit version 2.8.1 with options to split data into read pairs and in the original format (*ncbi/sra-tools*, 2008). Because *M. thibetana* was sequenced on an older platform, we used seqtk (Li, 2019) to convert from Q64 to Q33. Additionally, SRA files from multiple lanes as indicated in the FASTQ header were split for independent alignment to the reference genome facilitating the definition of separate read groups to be treated as independent in the variant calling workflow.

### Alignment, genome analysis, and variant calling

Raw FASTQ files from all samples were aligned to the reference genome rheMac8 (NCBI Accession: PRJNA214746). The masked reference genome was downloaded from UCSC. The reference genome for rheMac8 includes 20 autosomes, a pair of sex chromosomes, a mitochondrial genome, and 284,705 unnamed scaffolds of varying length. The total length of all scaffolds is 3.2Gb. The total length of all named chromosomes is 2.8 Gb. GATK best practices were followed to obtain high quality variant sites (see Supplementary Methods and Tables S1-S2). Briefly, reads were aligned to the reference genome, duplicates were marked, indels were realigned, base quality scores were recalibrated, variants were called, and variant quality scores were recalibrated. Samples from each species were processed independently and merged into a final callset (see Data Availability). The baboon reference genome was added to the variant files via two-way genome alignment and custom scripts that are available via GitHub. These final filtered files for each chromosome with the baboon information are freely available and were used for all subsequent analyses. A more detailed workflow is described in both the supplement as well as the GitHub page for the project, which includes command prompts and custom scripts (see Data Availability).

### Four-taxon test for introgression

To analyze patterns of introgression in sliding windows, a modified four-taxon test was used (Figure S2). Rather than separately analyzing gene flow between *M. arctoides* and each parental species group, as has been done previously (Fan *et al.*, 2018), we relied on the modified statistic *f_dM_* (Malinsky *et al.*, 2015), which is suggested to perform better than other related statistics based on simulation results (Martin *et al.*, 2015). Specifically, hybridization between the three samples of *M. arctoides* (P3) was tested with samples from the *sinica* species group (P1) and the *fascicularis* species group (P2), using the baboon reference genome as an outgroup (papAnu4; see Supplement for details). Our approach differed from the typical way these tests are setup where one may only be interested in shared ancestry between two pairs of taxa, (P1:P3 and P2:P3), and the P1 and P2 taxa are more closely related to each other than either are to P3 (see Figure 2 in Kulathinal *et al.*, 2009). However, this modification allowed for the quantification of shared ancestry from both putative parents together. It is worth noting that previous work had similar results when using the more traditional four taxon test setup for these taxa (Fan *et al.*, 2018).

Calculations of *f_dM_* were done using scripts available on GitHub (Martin, 2019) and using several differently sized genomic windows. *f_dM_* was computed for overlapping windows of size 5kb, 25kb, 50kb, 100kb, 500kb and 1000kb, with a step size of 20% of each window size. The minimum number of sites per window size was based on the distribution of sites available in the callset at each bin size. Specifically, a minimum number of sites of 10, 50, 100, 200, 1000, and 2000, respectively, per window size were used. Because it is recommended to have at least 100 sites per window (Martin, 2019), the 50kb window size was selected and the results are presented in Figure 2 as a heatmap across the genome. Results from the additional bin sizes are shown in Figure S3. This analysis was repeated with individual taxa from each species group (Table S3).

### Topology weighting

To analyze the weights of different phylogenetic topologies across the genome, the software Twisst was used to compare the weights of different species topologies throughout the genome (Martin & Belleghem, 2017; Martin *et al.*, 2019). First, data used for the introgression analysis were pruned to make sure that each group had at least one member represented in the genotype data. Next, neighbor joining trees were constructed in sliding windows of 50 SNPs each across the genome using Phyml from scripts available on GitHub (Martin, 2019). This generated newick formatted tree files that were used as input for Twisst. The resulting weight distributions among the three possible topologies were summed across the genome (Figure 2D).

The topology weighted outputs were further explored by collapsing adjacent intervals with the same majority topology supported. Since there were three possible topologies, the majority topology was defined as any one topology that had at least two-thirds of the total weight values. Any intervals without a majority topology were labeled as “unresolved”. If adjacent intervals supported the same majority topology, they were collapsed. The data were subsequently split into the three major topologies. These were intersected with gene regions from the Ensembl annotations for rheMac8 downloaded from UCSC.

Bootstrap confidence intervals for topo1, topo2, topo3, and unresolved proportions were computed by resampling 2,792 1 Mbp intervals along the genome (149 when considering only chromosome X) for each of 10,000 replicates and taking the 2.5% and 97.5% quantiles.

### Divergence Analysis

Based on the three possible topologies, nucleotide divergence was calculated as DXY using scripts available on GitHub (Martin, 2019). First, stringency in defining the majority topology was increased to require 100% of the weights be on only one of the three topologies. Next, divergence was calculated between *M. **arctoides and members of the fascicularis species group (M.** fascicularis* and *M. mulatta*) for regions with regions with topo3 ((arc,fas),sin). ***As well, divergence was calculated between M. arctoides and members of the sinica species group (*M. assamensis* and *M. thibetana*) for genomic regions with topo2 (fas,(arc,sin))***. For the regions with topo1 (arc,(fas,sin)), the relative divergences were unchanged when divergence was calculated between *M. arctoides* and members of the *sinica* species group as compared to divergence calculated between *M. arctoides* and members of the *fascicularis* species group, therefore only the latter is reported.

### Candidate Gene Selection

A major criterion of homoploid hybrid speciation is evidence that RI evolved as a by-product of hybridization (Schumer *et al.*, 2014). This coupled with the unique genital morphology of both males and females in the bear macaque (Figure 1) led us to focus our study on candidate genes that have been identified in other studies to be associated with genital morphology. We conducted a literature search for terms associated with genital morphology such as “disorder of sex development”, “DSD”, “baubellum”, and “baculum”, the latter two terms included due to the extremely large baculum of the bear macaque, and the female equivalent structure. We filtered to remove genes that solely associated with either fertility or adult hormonal disorders/shifts as these conditions are unlikely to affect the development of genitalia. Included in our initial list of 141 genes (Table S4) were genes associated with gonad dysgenesis (Bamshad *et al.*, 1997; Ekici *et al.*, 2013; Witchel, 2018; Lavi *et al.*, 2020), female masculinization (Wang *et al.*, 2020), genital tubercle formation (Haraguchi *et al.*, 2000), baculum morphology (Schultz *et al.*, 2016a), hypogonadism (Délot *et al.*, 2017), gonad differentiation (Witchel, 2018), gonad development (Witchel, 2018), sex determination (Délot *et al.*, 2017), and sex differentiation (Délot *et al.*, 2017).

In addition to our literature search, we used the Human Phenotype Ontology (HPO) (Robinson *et al.*, 2008) database to target search terms associated with genital abnormalities, specifically targeting those that would likely be developmental. We therefore targeted the HPO category “Abnormal Reproductive System Morphology”, but removing categories that were unrelated, such as “Genital Ulcers”. A list of the HPO terms can be found in Table 2. In total, this added 1230 genes to our 141 set of literature candidate genes. It is worth noting that 88 genes overlapped between these lists, with the remainder being unique to our literature search.

**Table 2.** Summary of candidate gene lists. In addition to a manually curated list from the literature, several phenotype terms were mined from the Human Phenotype Onotology (HPO) and the Mammalian Phenotype (MP) Databases. Here the number of genes associated with each term is listed along with the number of orthologs in rheMac8. Further, the number of genes used in our analysis where we calculated seven population genetics metrics are listed along with the final number of outliers per candidate gene set.

| Unique ID | Subontology | Database | # of Genes | # rheMac8 orthologs | # genes analyzed dN/dS | # genes analyzed other stats | # outliers |
| --- | --- | --- | --- | --- | --- | --- | --- |
| **n/a** | **Manually curated set** | **Literature** | **141** | **110** | **17** | **64** | **7** |
| HP:0000811 | Abnormal external genitalia | HPO | 1061 | 920 | 140 | 425 | 33 |
| HP:0010461 | Abnormality of the male genitalia | HPO | 1053 | 907 | 142 | 424 | 34 |
| HP:0010460 | Abnormality of the female genitalia | HPO | 512 | 447 | 72 | 223 | 15 |
| HP:0000812 | Abnormal internal genitalia | HPO | 468 | 427 | 62 | 196 | 14 |
| HP:0001827 | Genital tract atresia | HPO | 16 | 15 | 3 | 10 | 1 |
| HP:0012244 | Abnormal sex determination | HPO | 20 | 17 | 2 | 8 | 1 |
| HP:0003252 | Anteriorly displaced genitalia | HPO | 1 | 1 | 1 | 2 | 0 |
| MP:0009198 | abnormal male genitalia morphology | MP | 1145 | 959 | 130 | 478 | 24 |
| MP:0009208 | abnormal female genitalia morphology | MP | 696 | 587 | 89 | 304 | 18 |
| MP:0002210 | abnormal sex determination | MP | 782 | 644 | 90 | 315 | 19 |
| MP:0003936 | abnormal reproductive system development | MP | 99 | 91 | 12 | 46 | 1 |
| **Grand total** |  |  |  | **2284** | **313** | **1055** | **66** |

Additionally, because certain phenotypes are lacking in humans, such as the baculum, we did a similar search of the Mammalian Phenotype (MP) Browser (http://www.informatics.jax.org/vocab/mp_ontology) available through the Mouse Genome Database. We used largely overlapping search terms to the HPO terms, totaling a set of 1554 genes. As expected, there were several genes that overlapped across each set, but also many genes unique to each query. In sum, our final candidate gene list comprised of 2,284 genes from the literature, as well as the HPO and MP databases that had orthologs in the rheMac8 reference genome (Table 2). Of these 2,284 genes, only a subset was used in the calculations of each population genetic metric based on constraints of these subsequent analyses. The major limiting factor was the data for recombination rate, which was used to identify genomic outliers. The total number of genes analyzed for each statistic is reported in Table S6 and summarized in Table 2. For most statistics, 1,055 candidate genes were included in the population genetic analysis for outliers.

### Outlier Analysis

With the candidate gene list, we sought to identify if any genes on our list had extreme values for diversity, differentiation, or introgression within the bear macaque. To do this, we took a gene centered approach. Instead of the pre-defined windows used earlier, we used the Ensembl Gene Annotations for rheMac8 to define window coordinates. First, the smallest region containing all transcripts from the annotation file was computed. Then, 5kb upstream and downstream was added to include potential regulatory regions for each gene. These windows were used to re-calculate *f_dM_*, and to calculate nucleotide diversity, pi. We also chose to look at two metrics for nucleotide divergence – F_ST_ and D_XY_ – calculated to compare *M. arctoides* to members of the both the *fascicularis* and *sinica* species groups separately. All metrics were calculated using scripts available on GitHub (Martin, 2019). Finally, we estimated dN/dS for each gene. This methodology was previously described in (Bailey & Stevison, 2021). Briefly, we used the three possible gene tree topologies to estimate a branch specific dN/dS for the *M. arctoides* branch for every gene in the genome. Estimates where dS was 0 and dN/dS was greater than 10 were excluded as these genes had insufficient divergence to obtain valid estimates (Villaneuva-Canas *et al.* 2013). Then, the topology weights for each gene window from our Twisst analysis were used to weight the dN/dS estimates for each gene across the three gene trees.

To further identify extreme values of each metric, we took into consideration that many of these statistics have been shown to be correlated with recombination rate, which can impede the detection of outliers, specifically enriching for outliers in regions of low recombination rate (Stevison & McGaugh, 2020). Therefore, we also computed an average recombination rate for each gene. For this, we download the UCSC track for recombination rate associated with rheMac8 (ftp://ftp.hgsc.bcm.edu/ucscHub/rhesusSNVs/rheMac8/all.rate.bw) (Xue *et al.*, 2016) and converted it from bigwig to bedGraph format using UCSC utilities. Bedtools (Quinlan & Hall, 2010) was then used to calculate the average recombination rate (cM/Mb) in each gene interval.

Finally, the Cook’s distance (cooksd) (Cook & Weisberg, 1982) was calculated from a linear regression between each statistic and recombination rate per gene. Cook’s distance provides a measure of the leverage and residual values of a particular data point, which indicates how much it influences the overall regression. Thus, it reveals how much the regression changes when that observation is removed, making it useful for identifying outliers in a regression analysis. Here, extreme values were determined based on genes that were 3*mean(cooksd) above or below the fitted value in the regression model, depending on the particular statistic. The resulting outlier genes were genes where the statistic being evaluated (e.g. F_ST_, D_XY_) was much higher/lower than expected when considering the relationship between the statistic and recombination rate. For *f_dM_*, because it is a two-tailed distribution, outliers from both the upper and lower end of the distribution were computed. For dN/dS, we did not find a statistically significant relationship with recombination rate, consistent with prior findings in humans (Coop & Przeworski, 2007). Therefore, we used a 5% cutoff for extreme values of this statistic instead of the Cook’s distance approach. This percentage produced fewer genome-wide outliers than the average of all other metrics, suggesting it as an appropriate cutoff (Table S6).

Genes identified as genome-wide outliers were then compared against our set of a priori candidate genes to identify outliers that were putatively involved in genital morphology. A chi-square test was conducted to determine if the number of outliers that were in our candidate gene set were more than expected due to random chance. A summary of the regression results and chi-square tests can be found in Table S6, and the resulting 66 outlier genes that overlapped our candidate gene set can be found in Table S7. BCFtools (version 1.11) (Li *et al.*, 2009a) view was used to extract VCFs for the 66 outlier genes and these 66 VCFs were combined and annotated using SnpEff (version 4.3p) (Cingolani *et al.*, 2012) to examine the putative impacts of variants identified within this system. The Mmul_8.0.1.86 annotation was downloaded through SnpEff to conduct this analysis as this corresponds to the reference genome used for samples in this study. These annotations include exons, introns, UTRs and regions 5kb upstream and downstream of start and end sites for genes to account for possible regulatory factors. These annotations are provided in tabular format in Table S7.

### Permutation Analyses

Permutation tests were used to identify differences in various summary statistics (F_ST_, D_XY_, *f_dM_*, dN/dS, and pi) between candidate genital genes and all other genes. For calculations of pi this was a one-tailed test where genital genes were expected to have reduced pi in *M. arctoides* compared to other genes. For calculations of dN/dS this was a one-tailed test where genital genes were expected to have increased dN/dS in *M. arctoides* compared to other genes. All other comparisons (F_ST_, D_XY_, and *f_dM_*) were two-tailed tests. In all of these tests the test statistic used was the difference of the mean summary statistic for the candidate genital genes and the mean for the same summary statistic for all other genes. These analyses were conducted using modified perl scripts from Evans *et al.* (2021) and the results are reported in Table S6.

In addition to this, we conducted permutation analyses on genes categorized as being involved in either male or female genital development or abnormalities according to the HPO and MP databases. Specifically, the categories used were HP:0010461, HP:0010460, MP:0009198 and MP:0009208 (see Table S5). This was done to examine the possibility of male and female genital morphology evolving at different rates (see Introduction). These analyses parallel the above in examining the same summary statistics, though all tests were two-tailed as there was not a specific expectation for the direction of evolution.

## Results

Genomic analysis of five macaque species revealed a mosaic of evolutionary ancestry with respect to two major species groups. These analyses used newly sequenced whole genome samples from *M. arctoides* (20X coverage) and *M. assamensis* (13X), and 8 publicly available genomes ranging in coverage from 4-49X (median 32X; Table 1). Most variable positions were heterozygous in only one species (Figure 2C). For shared variants, *M. arctoides* shares the most variants with the *sinica* species group, *M. assamensis* and *M. thibetana*, consistent with taxonomic placement in this species group (Fooden, 1980). However, the two-way intersection between *M. arctoides* and *M. thibetana* had the fewest shared heterozygous sites, which is likely due to the low overall heterozygosity in the *M. thibetana* sample (Figure 2C; Table 1). *M. assamensis* had the largest number of heterozygous sites (∼4.7 million; Table 1; Figure 2C).

### Extensive Mosaicism of the *M. arctoides* Genome

The four-taxon test (Kulathinal *et al.*, 2009; Green *et al.*, 2010; Martin *et al.*, 2015) results which used all three *M. arctoides* samples, supported that the *M. arctoides* genome has a mosaic of shared ancestry with both the *sinica* and *fascicularis* species groups (Figure 3), which is consistent with extensive introgression from both species groups, though some of these windows with shared ancestry could also be due to incomplete lineage sorting between these taxa (Stevison & Kohn, 2009) (See Supplementary Materials). The genome-wide average *f_dM_* (Malinsky *et al.*, 2015; Martin, 2019) value was −0.113, ranging from −0.69 (with negative values supporting shared ancestry with the *sinica* group) and 0.55 (with positive values supporting shared ancestry with the *fascicularis* group) in 50kb sliding windows. This sliding window analysis was repeated with several window sizes (Figure S3), and all had a similar genome-wide mean *f_dM_*. In a previous study, estimates of Patterson’s D (Green *et al.*, 2010) were similar to the *f_dM_* estimates here (see Table S8 in (Fan *et al.*, 2018)).

**Figure 3.**
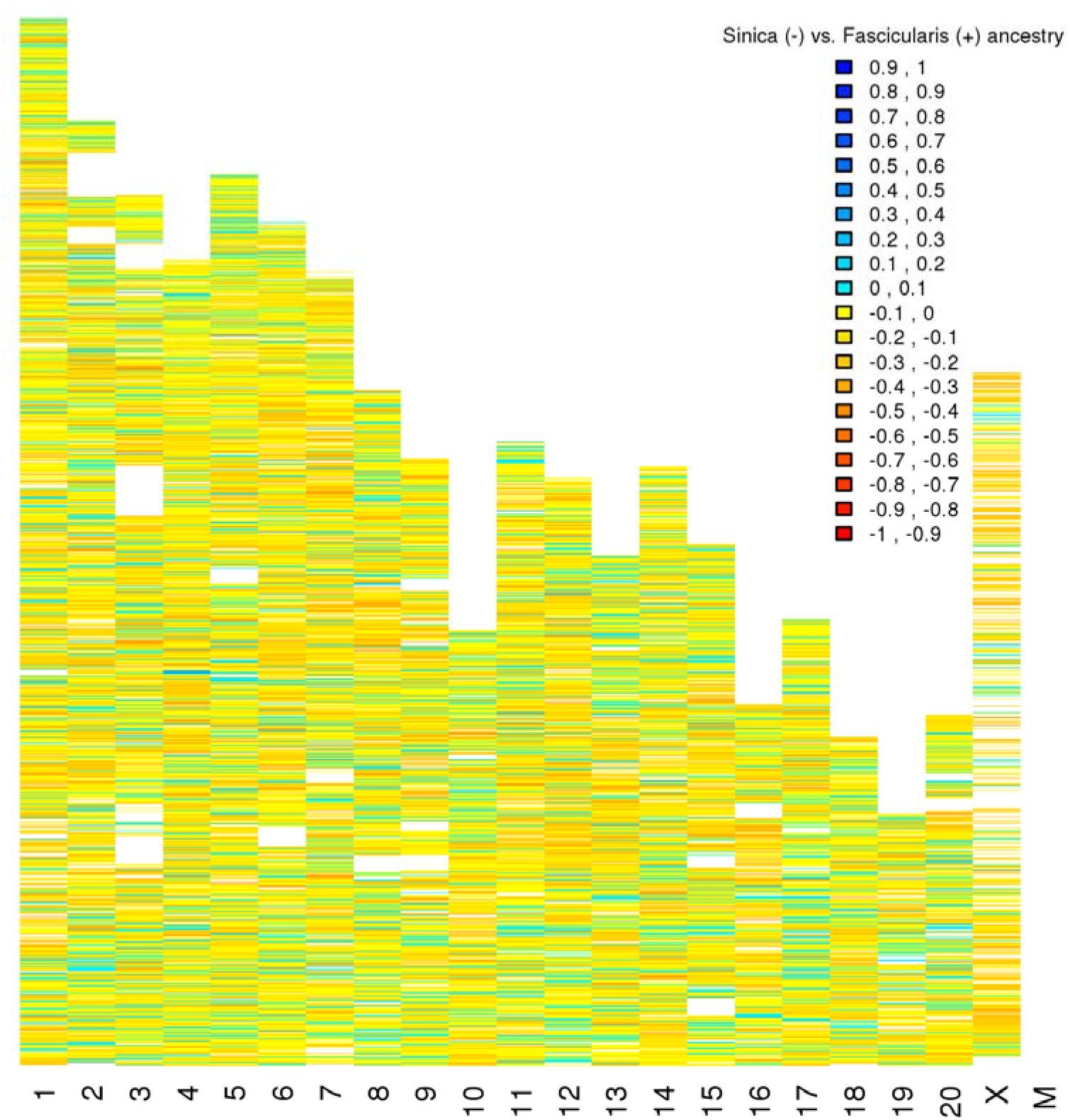
Introgression analysis. Results from analysis of introgression (*f_dM_*) between *M. arctoides* samples and the parent taxa in 50kb sliding windows. Regions where *M. arctoides* has most recent ancestry with the *sinica* group are displayed as negative values and regions where *M. arctoides* shares most recent common ancestry with the *fascicularis* group are displayed as positive values. Other window size results are in Figure S3 and plots using individual *M. arctoides* samples are in Figure S4.

Phylogenetic relationships among the species groups were evaluated via the software Twisst (Martin & Belleghem, 2017) (topology weighting by iterative sampling of subtrees). This method quantifies phylogenetic relationships across the genome and returns weights in sliding windows (Figure 2D). The topology weights were used to define a “majority topology” as having two-thirds or more of the sum of weights in any region. Windows that did not meet this criterion were re-classified as “unresolved”. This analysis showed the proportion of the genome that supported “topo2” (fas,(arc,sin)), which groups *M. arctoides* with the *sinica* group, as 52.64% (95% bootstrap CI [52.02%, 53.28%]). However, a significant portion, 15.70% (95% bootstrap CI [15.40%, 16.01%]), of the genome supported “topo3” ((arc,fas),sin), which groups *M. arctoides* with the *fascicularis* species group. Interestingly, 11.07% (95% bootstrap CI [10.84%, 11.31%]) of the genome supported *M. arctoides* clustering in a group by itself outside of the *sinica* and *fascicularis* groups (“topo1”; (arc,(fas,sin))). The remaining 20.59% (95% bootstrap CI [20.18%, 21.01%]) of the genome was categorized as unresolved because none of the three topologies had >2/3^rds^ weight. That the 95% confidence intervals of topo3 do not overlap with those of topo1 is consistent with the putative hybrid origin of *M. arctoides*, as opposed to topo1 and topo3 both stemming from ancestral polymorphism.

However, a subsequent investigation of nucleotide divergence of these genomic regions revealed that D_XY_ between sister taxa in regions supporting topo2 (fas,(arc,sin)) were significantly lower than regions supporting topo3 ((arc,fas),sin) (Figure S5). This result is consistent with either *M. arctoides* being sister to the *fascicularis* group with subsequent hybridization with the *sinica* species group (Figure S1B), or that *M. arctoides* split from macaques prior to the split of the *sinica* and *fascicularis* species groups with subsequent hybridization from each group at different times (Figure S1C). This latter scenario is consistent with the deepest nucleotide divergence being in regions supporting topo1 (arc(sin,fas)). However, it is worth pointing out that this analysis does not include additional members of the *sinica* species group (e.g. *M. sinica* and *M. radiata*) that would be crucial to resolving this complex evolutionary history (see Supplementary Materials).

### Investigation of outlier genes associated with genital morphology

Our gene centered approach led to the calculation of seven major population genetic statistics across all genes in the rheMac8 annotation. In total, our analysis focused primarily on 10,543 genes for which we were able to get reliable estimates for each metric, including recombination rate (Table S6). However, for the dN/dS calculations, this baseline number of genes was far fewer with only 1903 due to the limitation of our approach (see Methods). Of the 2,284 genes that were included on our candidate gene list (Table 2 and S4) and also had orthologs in rheMac8, only 1,055 were included in our most analyses (Table 2 and S6). Of the genome-wide outliers across these metrics, 66 of these 1,055 genes were statistically elevated as potential outliers for further investigation (Table S7).

A chi-square test was used to compare the proportion of outlier candidate genes (OCGs) against those analyzed to the expected number of outliers based on the remainder of the genes in the genome (Table S6). This analysis revealed that there were no more outliers than expected by chance, suggesting that the 66 outliers is not more than would be expected of any other set of candidate genes. However, a permutation test was used to compare the difference between means across the seven population genetic metrics found that for four metrics, the mean of the candidate genes was significantly different than expected based on the value across the remainder of the genes in the genome (Figure 4). These metrics showed that the nucleotide diversity, pi, across the candidate genes was significantly lower than expected (Figure 4A). Additionally, both measures of F_ST_ between *M. arctoides* and the two species groups revealed higher than expected nucleotide divergence (Figure 4C-D). Similarly, D_XY_ between *M. arctoides* and the *fascicularis* species group was much higher than expected based on 100 permutations of a similar number of genes that were not part of the candidate gene set.

**Figure 4.**
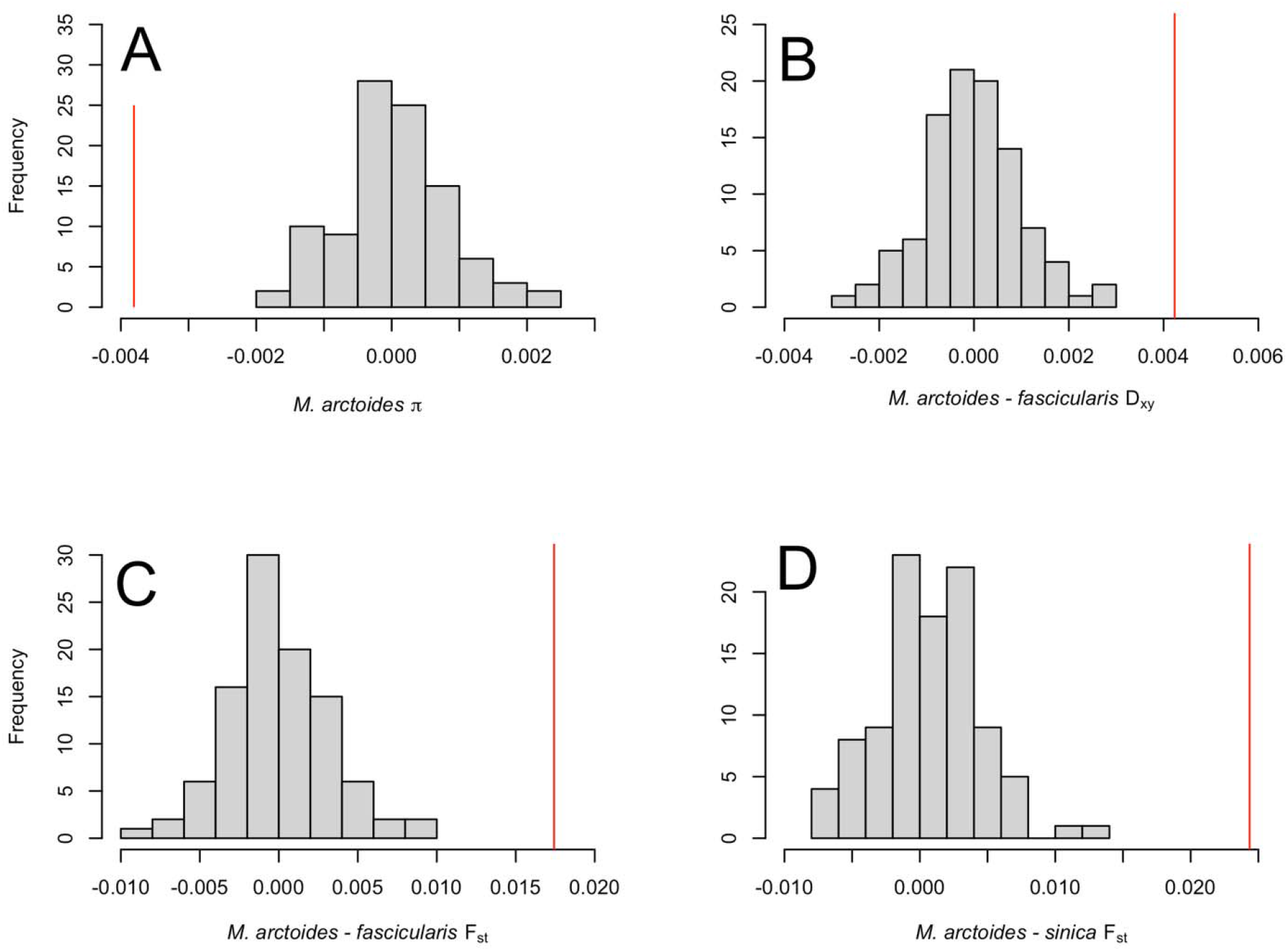
Permutation test results. For each panel, the red line is the difference of the mean ogenital genes and all other genes for a given summary statistic. Additionally, a histogram of the results from 100 permutations where the mean difference was recalculated using the same numbe of randomly selected genes as the candidate genital genes is shown. The various summary statistic plotted here are A) pi in *M. arctoides*, B) D_XY_ between *M. arctoides* and *fascicularis* group species, C F_ST_ between *M. arctoides* and *fascicularis* group species, and D) F_ST_ between *M. arctoides* and sinic group species. The computed p-values for all tests shown here were zero. The remaining statistic were not significantly different and are reported in Table S6 and Figure SX.

An investigation into the 66 genes found that three genes were OCGs across four statistics (CEP19, GATA3, and PTPN23). Three more genes were OCGs across three statistics (AIRE, CSPP1, and SNAI2). Finally, 11 genes were OCGs across two of the seven statistics (CAMKV, DND1, HMGA1, HSPA4, KDM3B, PIK3CA, PSMB8, SIN3A, TDRKH, TMED10, and ZCWPW1). It is worth noting that although the genes on the HPO and MP database lists were under the specific search terms, each of these genes were associated with many unrelated phenotypes, thus it is difficult to interpret their overall significance in this analysis. For OCGs from the HPO database, each gene had an average of 70 phenotypic terms with a range from 4 to 302. For the OCGs in the mammalian phenotype database, the average number of phenotypic terms was smaller at 22.6, and the range was from 1 to 96. This suggests the mammalian database may be slightly better curated to avoid genes that are overly pleiotropic. Interestingly, of these 17 OCGs that were outliers across more than one statistic, only two have putative high impact variants (CEP19 and CSPP1), with the latter also having 15 missense mutations of putative moderate impact. CSPP1 is associated with 151 HPO terms, being presumably involved in a lot of different phenotypes (Table S7).

Despite the uncertainty of hits in the larger databases, seven of the OCGs were on the original literature survey for candidate genes (Table 2 and S4), which were more carefully selected based on their role in genital morphology. Of these, GATA3 was an outlier across multiple statistics, which was initially included due to its known associated with urogenital anomalies (Délot *et al.*, 2017), with over 20 years of research on this gene (Lemos & Thakker, 2020), including knockout studies (Grote *et al.*, 2008). Another of these seven is HFE, which is associated with hereditary hemochromatosis (HH) and subsequent hypogonadism (Délot *et al.*, 2017) due to decreased iron circulation (Bezwoda *et al.*, 1977; McDermott & Walsh, 2005), with symptoms such as decreased libido and testicular atrophy in humans (Kelly *et al.*, 1984). In our SNP effect analysis, GATA3 had 1 predicted moderate impact variant and 8 low impact variants (Table S7).

Additionally, two of the seven OCGs from the literature survey gene list were CYP proteins (CYP17A1 and CYP21A2). Both CYP17A1 and CYP21A2 are associated with congenital adrenal hyperplasia (CAH aka 17OHD), a disorder which leads to abnormal sexual development (Délot *et al.*, 2017), with CYP21A2 mutations being the most common cause of CAH (Koppens *et al.*, 1992). CYP21A2 mutations in humans have been associated with simple virilization, in which female genitalia are masculinized during development (Koppens *et al.*, 1992). Several cohort studies have found genotype-to-phenotype associations linking CYP21A2 mutations to a variety of CAH phenotypes (Torres *et al.*, 2003; Soardi *et al.*, 2008; de Carvalho *et al.*, 2016). Additionally, protein structure analysis has been done to examine the impact of specific mutants on a form of CAH known as salt-water wasting disease, which if left undiagnosed leads to low sodium levels and death (Haider *et al.*, 2013). CYP17A1 is heavily involved in steroidogenesis, with deletion mutations leading to hormone production inhibition and sex development abnormalities (Keskin *et al.*, 2015; Turkkahraman *et al.*, 2015; Zhang *et al.*, 2015; Papi *et al.*, 2018; Xia *et al.*, 2021). Recent genome studies of 17OHD patients revealed 46,XY karyotype in physically presenting females which co-occur with multiple mutations in the CYP17A1 gene (Zhang *et al.*, 2015; Xia *et al.*, 2021). CYP21A1 had two high, 21 moderate, and 32 low impact variants, whereas CYP17A1 had two moderate and eleven low impact variants (Table S7). Additionally, CYP21A1 had the highest proportion of silent and nonsense variants, as well as the 2^nd^ highest proportion of missense variants (Figure 5).

**Figure 5.**
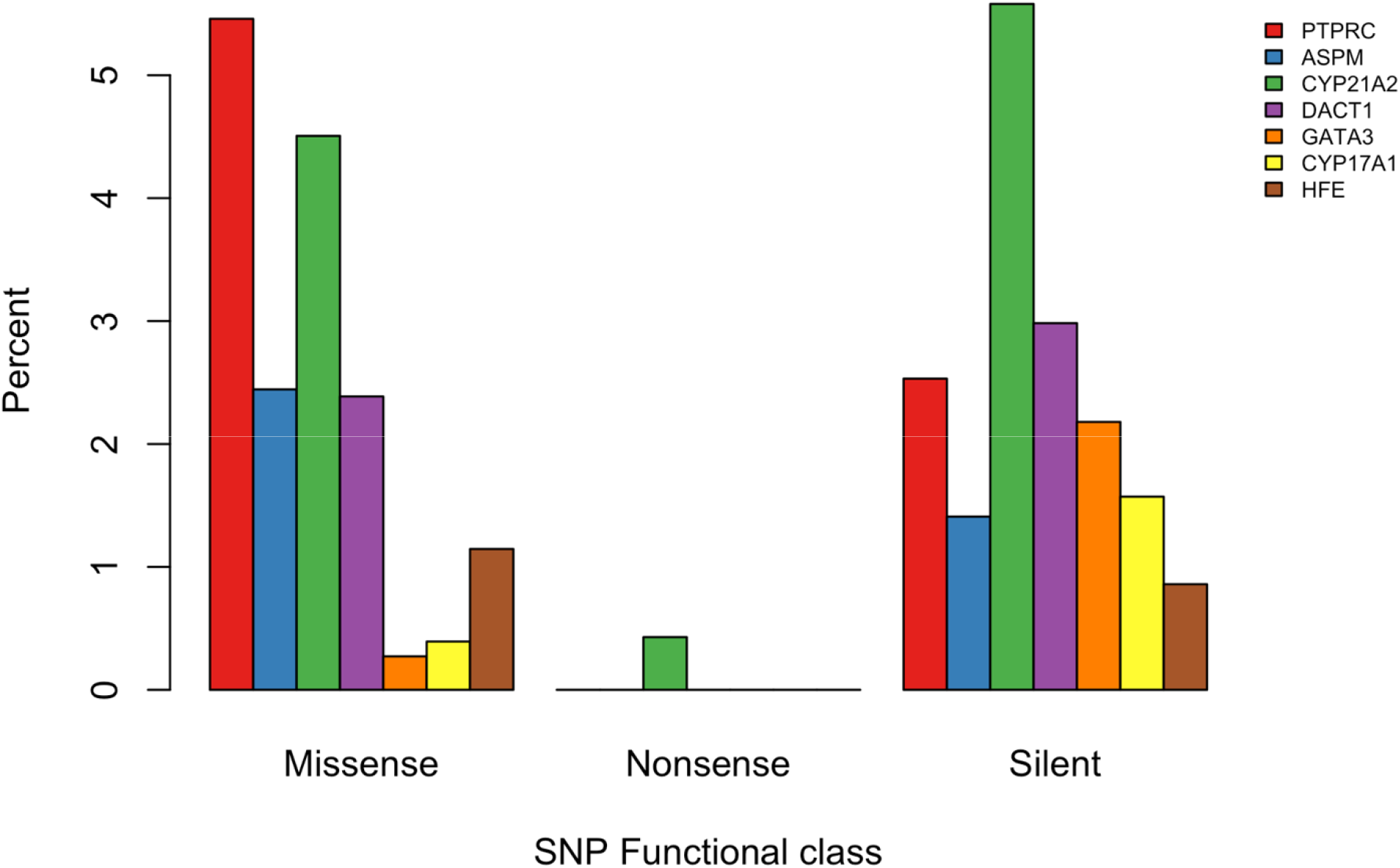
Functional Impact of Variants in Outlier Candidate Genes. Results from SNPeff are shown broken down by functional impact to the protein sequence of the seven outlier candidat genes based on the literature list. Functional classes are missense variants which change the amin acid, nonsense, which result in a premature stop codon, and silent variants, which alter the nucleotide sequence, but result in a synonymous amino acid.

Three more OCGs on the original literature candidate gene list that were outliers for dN/dS were identified in a study as major candidates for baculum morphology in mice (Schultz *et al.*, 2016a) (DACT1, PTPRC, and ASPM; Figure 4). This is particularly intriguing when one considers the unique baculum morphology of the os penis in the bear macaque relative to all other primates (see Discussion). Because these were outliers for dN/dS, which is targeted to the protein coding regions of the gene, we further investigated the protein sequence of each gene to look at coding mutations. PTPRC had the highest proportion of missense mutation among the seven literature outliers (Figure 5). Specifically, there were seven amino acid changes in multiple individuals, two of which overlapped the fibronectin domain and had significant changes in the amino acid properties. Two of these changes were completely fixed in *M. arctoides* (p.K449E and p.R424R), two were nearly fixed, shared only with a single *M. assamensis* sample (p.E314G and p.Y359S), and three amino acid changes were shared between *M. arctoides* and members of the *fascicularis* species group (p.D363G, p.K384N, and p.D385N). DACT1 had the 2^nd^ highest proportion of silent variants among the seven literature OCGs (Figure 5). Additionally, we found 3 major amino acid changes clustered together, two that were nearly fixed in *M. arctoides* as compared to all other macaques (p.C439Y and p.G493D). Additionally, there was a p.Q498R mutation that was shared between *M. arctoides* and members of the *fascicularis* species group. This similarity is of particular interest due to the shared baculum morphology of these species (see Discussion). Finally, for ASPM, there was only one fixed amino acid change (p.H352P), which overlapped a region of the protein that interacts with another enzyme called katanin. Additionally, there was a p.H1870R mutation that was shared with *sinica* species, and several other amino acid changes that were unique to *M. arctoides*, but polymorphic rather than fixed (p.V269F, p.T357I, and p.R1164H). While all of these amino acid changes are likely to have an impact on the protein, the specific changes to the gene function and its overall implications for the baculum phenotype are unknown. Still, these residues would be of particular interest in mouse mutant studies on the genetics of baculum morphology.

Permutation analysis of male versus female genital development genes showed no significant difference in all statistics except *f_dM_* (p = 0.02), where male genital genes exhibited higher introgression between *M. arctoides* and *sinica* species than female genes (Table S5).

## Discussion

Genital morphology genes have higher divergence and lower diversity than expected Although the number of outliers from our genital candidate genes was not more than expected by chance, the average divergence was higher than expected and the diversity was lower than expected (Figure 4). In fact, none of the permutations overlapped the observed mean difference between the actual candidate gene set for four of the seven population genetic metrics. Specifically, the candidate genes had lower nucleotide diversity (pi) which is consistent with recent positive selection on these genes that would have reduced local genetic variation. Similarly, F_ST_ between *M. arctoides* and both species groups was higher than expected, which is also consistent with positive selection on this subset of genes as compared to the remainder of the genome. Additionally, D_XY_ between *M. arctoides* and *fascicularis* species was elevated, but not D_XY_ between *M. arctoides* and *sinica* species. These results suggest that the evolution of these genes have been increased relative to the rest of the genes in the genome, confirming the importance of genital morphology in the evolution of this species. A possible explanation for the difference in the two measures of nucleotide divergence is that F_ST_ is a relative measure of divergence, whereas D_XY_ is an absolute measure (Noor & Bennett, 2009). Here, the majority of the genome clusters with *sinica* species, which could explain why D_XY_ between *M. arctoides* and the *sinica* species group does not have more outliers than expected.

Of 66 candidate genes identified as outliers, many are involved in a variety of phenotypes. Of course, because these are developmental phenotypes, it is not surprising that the impact of these genes would be highly pleiotropic. However, their specific role in contributing to the divergent genital morphology and subsequent reproductive isolation in this species is unclear. One interesting thing worth noting is that the number of genes in both phenotype onotology databases associated with male genital abnormalities is nearly 2x the number of genes associated with female genital abnormalities (Table 2). This is not likely due to underlying differences in the genetics of these traits, but instead more likely reflects a bias towards research into male reproductive morphology as compared to female reproductive abnormalities. Still, for the most part, the mean values across our various metrics were not significantly different between these sets of male vs. female genes (Table S5). While the result for *f_dM_* was statistically significant, both values were lower than the genome-wide mean, and thus do not likely reflect differences in the direction of introgression. In contrast, the difference in means of dN/dS, though not statistically significant, had twice the difference of *f_dM_*. Specifically, the mean dN/dS across male genes (0.141) matches the mean of the genome background (0.143), whereas the mean across female genes was much lower (0.101). Though speculative, this difference is perhaps consistent with the idea that many genes of large effect on female genital morphology have yet to be described, supported by the disparity in the gene set sizes. Together these findings suggests that future work should aim to identify additional potentially undiscovered genes that play a role in female genital morphology.

### Divergent baculum morphology putatively explained by dN/dS outliers

Regardless of the inconclusiveness of the genes that were found based on phenotype ontology methods, 3/21 outliers for dN/dS were genes previously identified in mice as candidate genes for morphological variation in the baculum (DACT1, PTPRC, and ASPM). The baculum, or penis bone, is a novel male genital structure in placental mammals that evolved ∼145 million years ago (Brindle & Opie, 2016), with the bear macaque, having the longest primate baculum. Since that time, it has been gained or lost multiple times, including recent loss in Homo sapiens (Schultz *et al.*, 2016b; Carosi & Scalici, 2017). Despite its remarkable interspecific variation, the penis bone generally does not vary extensively within species (Ramm *et al.*, 2009), and may be the only physical trait that differs consistently between closely related species, making it a key morphological character for species identification (Carosi & Scalici, 2017). However, the genetics of morphological variation in this trait are only beginning to be understood. In 2016, a morphometric analysis using micro-CT scans of bacula in mice identified three major quantitative trait loci (QTLs), two that together explained 50.6% of the variation in baculum size, and one that explained 23.4% of the variation in baculum shape (Schultz *et al.*, 2016a). These regions were further narrowed by both differential gene expression in early development of male mice, and potential involvement in bone or genital morphogenesis, revealing 16 major candidate loci with potential effects on baculum development and morphology (see Table S3 in (Schultz *et al.*, 2016a)), that were included on our candidate gene list (Table S4).

The baculum in *M. arctoides* is more than double the average length of *sinica* species group taxa and nearly four times as long as *fascicularis* species group taxa (Fooden, 1971; Dixson, 1987). In addition to its elongation, the morphology of the *M. arctoides* baculum also has a unique arced crescent shape, with an absent distal region that is present in most other macaque species (See comparison in Figure 8 of Fooden *et al.*, 1985; Fooden, 1990). Among macaques, these two features – a gentle dorsoventral curvature and the lack of a well-defined distal process – are shared only by members of the *fascicularis* species group. This latter point perhaps explains why 4/5 amino acid changes across these three genes that were shared between *M. arctoides* and another group, were shared between *M. arctoides* and the *fascicularis* species members. This is in stark contrast to the rest of the genome which has a much higher proportion of shared variation with *sinica* species group members.

### Future Directions

This study has focused on genes with phenotypic associations with genital morphology, but an outlier analysis can only present hypotheses of the specific genes involved in the divergent reproductive traits of the bear macaque. Therefore, the 66 outlier genes are candidates that may have contributed to speciation. The target of future functional work should be to identify their specific role (if any) in contributing to either divergent male genital morphology or compensatory changes in female reproductive morphology. They should also be the target of functional studies in model systems, such as mice, on genital morphology of both males and females. For example, the seven fixed amino acid changes in the bear macaque across the three dN/dS outlier genes associated with mouse baculum morphology should be prioritized in future work on the genetics of this trait. Still, why this remarkable trait is divergent in the bear macaque is unknown. Perhaps more detailed functional connections between specific genes/variants and the baculum would allow for future work to examine the selective processes that shaped this unique genital phenotype.

Additionally, as the genetic basis of other genital traits, such as baubellum morphology become known, the candidate gene list can be extended. Similarly, as the genome annotation improves with subsequent reference genomes, more genes can be considered that have clear orthologs in macaques.

In addition to this study, there has been extensive interest in the evolution of this species. However, due to the limited number of publicly available whole genome samples (now three with the present study), the conservation status of this species, and the ethical and logistical barriers to experimental research with primates, it is challenging to make functional insights into its evolution. Additional WGS data from more individuals of this species would allow for a better characterization of selection/adaptation within this lineage. Additionally, more WGS samples would be useful in examining the distribution of introgression haplotype lengths to pinpoint the timing of events in this system that would better clarify when hybridization from each parent took place.

## Supporting information

Supplementary Methods and Results

## References

1. Bailey, N.P. & Stevison, L.S. 2021. Mitonuclear conflict in a macaque species exhibiting phylogenomic discordance. Journal of Evolutionary Biology 34: 1568–1579.

2. Bamshad, M., Lin, R.C., Law, D.J., Watkins, W.S., Krakowiak, P.A., Moore, M.E., et al. 1997. Mutations in human TBX3 alter limb, apocrine and genital development in ulnar-mammary syndrome. Nat Genet 16: 311–315.

3. Barnard, A.A., Fincke, O.M., McPeek, M.A. & Masly, J.P. 2017. Mechanical and tactile incompatibilities cause reproductive isolation between two young damselfly species: MATING STRUCTURES AND REPRODUCTIVE ISOLATION. Evolution 71: 2410–2427.

4. Bezwoda, W.R., Bothwell, T.H., Van Der Walt, L.A., Kronheim, S. & Pimstone†, B.L. 1977. An Investigation into Gonadal Dysfunction in Patients with Idiopathic Haemochromatosis. Clinical Endocrinology 6: 377–385.

5. Brindle, M. & Opie, C. 2016. Postcopulatory sexual selection influences baculum evolution in primates and carnivores. Proceedings of the Royal Society B: Biological Sciences 283: 20161736. Royal Society.

6. Carosi, M. & Scalici, M. 2017. Baculum (Os Penis). In: The International Encyclopedia of Primatology, pp. 1–5. American Cancer Society.

7. Cingolani, P., Platts, A., Wang, L.L., Coon, M., Nguyen, T., Wang, L., et al. 2012. A program for annotating and predicting the effects of single nucleotide polymorphisms, SnpEff: SNPs in the genome of Drosophila melanogaster strain w^1118^; iso-2; iso-3. Fly 6: 80–92.

8. Conway, J. & Gehlenborg, N. 2019. UpSetR: A More Scalable Alternative to Venn and Euler Diagrams for Visualizing Intersecting Sets. CRAN, https://CRAN.R-project.org/package=UpSetR.

9. Cook, R.D. & Weisberg, S. 1982. Residuals and Influence in Regression. New York: Chapman and Hall. Coop, G. & Przeworski, M. 2007. An evolutionary view of human recombination. Nat. Rev. Genet. 8: 23–34.

10. Danecek, P., Auton, A., Abecasis, G., Albers, C.A., Banks, E., DePristo, M.A., et al. 2011. The variant call format and VCFtools. Bioinformatics 27: 2156–8.

11. de Carvalho, D.F., Miranda, M.C., Gomes, L.G., Madureira, G., Marcondes, J.A.M., Billerbeck, A.E.C., et al. 2016. Molecular CYP21A2 diagnosis in 480 Brazilian patients with congenital adrenal hyperplasia before newborn screening introduction. Eur J Endocrinol 175: 107–116.

12. Délot, E.C., Papp, J.C., DSD-TRN Genetics Workgroup, Sandberg, D.E. & Vilain, E. 2017. Genetics of Disorders of Sex Development: The DSD-TRN Experience. Endocrinol Metab Clin North Am 46: 519–537.

13. Dixson, A.F. 1987. Observations on the evolution of the genitalia and copulatory behaviour in male primates. Journal of Zoology 213: 423–443.

14. Dixson, A.F. 1998. Primate sexuality: comparative studies of the prosimians, monkeys, apes, and human beings. Oxford University Press, Oxford; New York.

15. Dixson, A.F. & Anderson, M.J. 2004. Sexual behavior, reproductive physiology and sperm competition in male mammals. Physiology & Behavior 83: 361–371.

16. Eberhard, W.G. 1993. Evaluating Models of Sexual Selection: Genitalia as a Test Case. The American Naturalist 142: 564–571. The University of Chicago Press.

17. Eberhard, W.G. 2010. Evolution of genitalia: theories, evidence, and new directions. Genetica 138: 5–18.

18. Eberhard, W.G. 1996. Female control: sexual selection by cryptic female choice. Princeton University Press, Princeton, N.J.

19. Eberhard, W.G. 1985. Sexual selection and animal genitalia. Harvard University Press, Cambridge, Mass.

20. Ekici, A.B., Strissel, P.L., Oppelt, P.G., Renner, S.P., Brucker, S., Beckmann, M.W., et al. 2013. HOXA10 and HOXA13 sequence variations in human female genital malformations including congenital absence of the uterus and vagina. Gene 518: 267–272.

21. Elgvin, T.O., Trier, C.N., Tørresen, O.K., Hagen, I.J., Lien, S., Nederbragt, A.J., et al. 2017. The genomic mosaicism of hybrid speciation. Science Advances 3: e1602996.

22. Evans, B.J., Peter, B.M., Melnick, D.J., Andayani, N., Supriatna, J., Zhu, J., et al. 2021. Mitonuclear interactions and introgression genomics of macaque monkeys ( Macaca) highlight the influence of behaviour on genome evolution. Proc. R. Soc. B. 288: 20211756.

23. Fan, Z., Zhao, G., Li, P., Osada, N., Xing, J., Yi, Y., et al. 2014. Whole-Genome Sequencing of Tibetan Macaque (Macaca thibetana) Provides New Insight into the Macaque Evolutionary History. Mol Biol Evol 31: 1475–1489.

24. Fan, Z., Zhou, A., Osada, N., Yu, J., Jiang, J., Li, P., et al. 2018. Ancient hybridization and admixture in macaques (genus Macaca) inferred from whole genome sequences. Molecular Phylogenetics and Evolution 127: 376–386.

25. Fooden, J. 1980. Classification and Distribution of Living Macaques (Macaca Lacepede, 1799). In: The Macaques: Studies in Ecology, Behavior, and Evolution (D. G. Lindburg, ed), pp. 1–9. Van Nostrand Reinhold Company, New York.

26. Fooden, J. 1967. Complementary Specialization of Male and Female Reproductive Structures in the Bear Macaque, *Macaca arctoides*. Nature 214: 939–941. Nature Publishing Group.

27. Fooden, J. 1971. Male external genitalia and systematic relationships of the Japanese macaque (Macaca fuscata Blyth, 1875). Primates 12: 305–311.

28. Fooden, J. 1990. The bear macaque,*Macaca arctoides*: a systematic review. Journal of Human Evolution 19: 607–686.

29. Fooden, J., Guoqiang, Q., Zongren, W. & Yingxiang, W. 1985. The stumptail macaques of China. American Journal of Primatology 8: 11–30.

30. Gompert, Z., Fordyce, J.A., Forister, M.L., Shapiro, A.M. & Nice, C.C. 2006. Homoploid Hybrid Speciation in an Extreme Habitat. Science 314: 1923–1925.

31. Green, R.E., Krause, J., Briggs, A.W., Maricic, T., Stenzel, U., Kircher, M., et al. 2010. A draft sequence of the Neandertal genome. Science 328: 710–722.

32. Greenway, R., McNemee, R., Okamoto, A., Plath, M., Arias-Rodriguez, L. & Tobler, M. 2019. Correlated divergence of female and male genitalia in replicated lineages with ongoing ecological speciation. Evolution 73: 1200–1212.

33. Grote, D., Boualia, S.K., Souabni, A., Merkel, C., Chi, X., Costantini, F., et al. 2008. Gata3 Acts Downstream of β-Catenin Signaling to Prevent Ectopic Metanephric Kidney Induction. PLOS Genetics 4: e1000316. Public Library of Science.

34. Haider, S., Islam, B., D’Atri, V., Sgobba, M., Poojari, C., Sun, L., et al. 2013. Structure–phenotype correlations of human CYP21A2 mutations in congenital adrenal hyperplasia. PNAS 110: 2605–2610. National Academy of Sciences.

35. Haraguchi, R., Suzuki, K., Murakami, R., Sakai, M., Kamikawa, M., Kengaku, M., et al. 2000. Molecular analysis of external genitalia formation: the role of fibroblast growth factor (Fgf) genes during genital tubercle formation. Development 127: 2471–2479.

36. Jiang, J., Yu, J., Li, J., Li, P., Fan, Z., Niu, L., et al. 2016. Mitochondrial Genome and Nuclear Markers Provide New Insight into the Evolutionary History of Macaques. PLoS One 11.

37. Kamimura, Y. & Mitsumoto, H. 2012. Lock-and-key structural isolation between sibling Drosophila species: Genital lock-and-key in Drosophila. Entomological Science 15: 197–201.

38. Kelly, T.M., Edwards, C.Q., Meikle, A.W. & Kushner, J.P. 1984. Hypogonadism in Hemochromatosis: Reversal with Iron Depletion. Ann Intern Med 101: 629–632. American College of Physicians.

39. Keskin, M., Uğurlu, A.K., Savaş-Erdeve, Ş., Sağsak, E., Akyüz, S.G., Çetinkaya, S., et al. 2015. 17α-Hydroylase/17,20-lyase deficiency related to P.Y27*(c.81C>A) mutation in CYP17A1 gene. J Pediatr Endocrinol Metab 28: 919–921.

40. Klaczko, J., Ingram, T. & Losos, J. 2015. Genitals evolve faster than other traits in Anolis lizards. Journal of Zoology 295: 44–48.

41. Koppens, P.F., Hoogenboezem, T., Halley, D.J., Barendse, C.A., Oostenbrink, A.J. & Degenhart, H.J. 1992. Family studies of the steroid 21-hydroxylase and complement C4 genes define 11 haplotypes in classical congenital adrenal hyperplasia in The Netherlands. Eur J Pediatr 151: 885–892.

42. Kulathinal, R.J., Stevison, L.S. & Noor, M.A.F. 2009. The Genomics of Speciation in Drosophila: Diversity, Divergence, and Introgression Estimated Using Low-Coverage Genome Sequencing. PLoS Genet 5.

43. Langerhans, R.B., Anderson, C.M. & Heinen-Kay, J.L. 2016. Causes and Consequences of Genital Evolution. Integr Comp Biol 56: 741–751. Oxford Academic.

44. Lavi, E., Zighan, M., Abu Libdeh, A., Klopstock, T., Weinberg-Shukron, A., Renbaum, P., et al. 2020. A Unique Presentation of XY Gonadal Dysgenesis in Frasier Syndrome due to WT1 Mutation and a Literature Review. Pediatr Endocrinol Rev 17: 302–307.

45. Leducq, J.-B., Nielly-Thibault, L., Charron, G., Eberlein, C., Verta, J.-P., Samani, P., et al. 2016. Speciation driven by hybridization and chromosomal plasticity in a wild yeast. Nature Microbiology 1: 15003.

46. Lemos, M.C. & Thakker, R.V. 2020. Hypoparathyroidism, deafness, and renal dysplasia syndrome: 20 Years after the identification of the first GATA3 mutations. Human Mutation 41: 1341– 1350.

47. Li, H. 2019. Toolkit for processing sequences in FASTA/Q formats: lh3/seqtk. Github, https://github.com/lh3/seqtk.

48. Li, H., Handsaker, B., Wysoker, A., Fennell, T., Ruan, J., Homer, N., et al. 2009a. The Sequence Alignment/Map format and SAMtools. Bioinformatics 25: 2078–2079.

49. Li, J., Fan, Z., Sun, T., Peng, C., Yue, B. & Li, J. 2018. Comparative Genome-Wide Survey of Single Nucleotide Variation Uncovers the Genetic Diversity and Potential Biomedical Applications among Six Macaca Species. Int J Mol Sci 19.

50. Li, J., Han, K., Xing, J., Kim, H.-S., Rogers, J., Ryder, O.A., et al. 2009b. Phylogeny of the macaques (Cercopithecidae: Macaca) based on Alu elements. Gene 448: 242–249.

51. Malinsky, M., Challis, R.J., Tyers, A.M., Schiffels, S., Terai, Y., Ngatunga, B.P., et al. 2015. Genomic islands of speciation separate cichlid ecomorphs in an East African crater lake. Science 350: 1493–1498.

52. Mallet, J. 2007. Hybrid speciation. Nature 446: 279–283.

53. Martin, O.Y. & Hosken, D.J. 2003. The evolution of reproductive isolation through sexual conflict. Nature 423: 979–982.

54. Martin, S. 2019. General tools for genomic analyses. Github, https://github.com/simonhmartin/genomics_general.

55. Martin, S.H. & Belleghem, S.M.V. 2017. Exploring Evolutionary Relationships Across the Genome Using Topology Weighting. Genetics 206: 429–438.

56. Martin, S.H., Davey, J.W. & Jiggins, C.D. 2015. Evaluating the Use of ABBA–BABA Statistics to Locate Introgressed Loci. Mol Biol Evol 32: 244–257.

57. Martin, S.H., Davey, J.W., Salazar, C. & Jiggins, C.D. 2019. Recombination rate variation shapes barriers to introgression across butterfly genomes. PLOS Biology 17: e2006288.

58. Mavárez, J. & Linares, M. 2008. Homoploid hybrid speciation in animals. Molecular Ecology 17: 4181–4185.

59. McDermott, J.H. & Walsh, C.H. 2005. Hypogonadism in Hereditary Hemochromatosis. The Journal of Clinical Endocrinology & Metabolism 90: 2451–2455.

60. *ncbi/sra-tools*. 2008. NCBI - National Center for Biotechnology Information/NLM/NIH.

61. Noor, M.A.F. & Bennett, S.M. 2009. Islands of speciation or mirages in the desert? Examining the role of restricted recombination in maintaining species. Heredity 103: 439–444.

62. Papi, G., Paragliola, R.M., Concolino, P., Di Donato, C., Pontecorvi, A. & Corsello, S.M. 2018. 46,XY Disorder of Sex Development Caused by 17α-Hydroxylase/17,20-Lyase Deficiency due to Homozygous Mutation of CYP17A1 Gene: Consequences of Late Diagnosis. Case Rep Endocrinol 2018: 2086861.

63. Quinlan, A.R. & Hall, I.M. 2010. BEDTools: a flexible suite of utilities for comparing genomic features. Bioinformatics 26: 841–2.

64. Ramm, S.A., Khoo, L. & Stockley, P. 2009. Sexual selection and the rodent baculum: an intraspecific study in the house mouse (Mus musculus domesticus). Genetica 138: 129.

65. Rieseberg, L.H. 2003. Major Ecological Transitions in Wild Sunflowers Facilitated by Hybridization. Science 301: 1211–1216.

66. Robinson, P.N., Köhler, S., Bauer, S., Seelow, D., Horn, D. & Mundlos, S. 2008. The Human Phenotype Ontology: a tool for annotating and analyzing human hereditary disease. Am J Hum Genet 83: 610–615.

67. Roos, C., Kothe, M., Alba, D.M., Delson, E. & Zinner, D. 2019. The radiation of macaques out of Africa: Evidence from mitogenome divergence times and the fossil record. Journal of Human Evolution 133: 114–132.

68. Schultz, N.G., Ingels, J., Hillhouse, A., Wardwell, K., Chang, P.L., Cheverud, J.M., et al. 2016a. The Genetic Basis of Baculum Size and Shape Variation in Mice. G3 (Bethesda) 6: 1141–1151.

69. Schultz, N.G., Lough-Stevens, M., Abreu, E., Orr, T. & Dean, M.D. 2016b. The Baculum was Gained and Lost Multiple Times during Mammalian Evolution. Integr Comp Biol 56: 644–656.

70. Schumer, M., Rosenthal, G.G. & Andolfatto, P. 2014. How Common Is Homoploid Hybrid Speciation? Evolution 68: 1553–1560.

71. Shively, C., Clarke, S., King, N., Schapiro, S. & Mitchell, G. 1982. Patterns of sexual behavior in male macaques. American Journal of Primatology 2: 373–384.

72. Simmons, L.W. & Fitzpatrick, J.L. 2019. Female genitalia can evolve more rapidly and divergently than male genitalia. Nat Commun 10: 1312.

73. Sloan, N.S. & Simmons, L.W. 2019. The evolution of female genitalia. J Evol Biol 32: 882–899.

74. Soardi, F.C., Barbaro, M., Lau, I.F., Lemos-Marini, S.H.V., Baptista, M.T.M., Guerra-Junior, G., et al. 2008. Inhibition of CYP21A2 Enzyme Activity Caused by Novel Missense Mutations Identified in Brazilian and Scandinavian Patients. The Journal of Clinical Endocrinology & Metabolism 93: 2416–2420.

75. Sota, T. & Kubota, K. 1998. Genital Lock-and-Key as a Selective Agent against Hybridization. Evolution 52: 1507–1513.

76. Stevison, L.S. & Kohn, M.H. 2009. Divergence population genetic analysis of hybridization between rhesus and cynomolgus macaques. Mol. Ecol. 18: 2457–2475.

77. Stevison, L.S. & McGaugh, S.E. 2020. It’s time to stop sweeping recombination rate under the genome scan rug. Molecular ecology.

78. Torres, N., Mello, M.P., Germano, C.M.R., Elias, L.L.K., Moreira, A.C. & Castro, M. 2003. Phenotype and genotype correlation of the microconversion from the CYP21A1P to the CYP21A2 gene in congenital adrenal hyperplasia. Braz J Med Biol Res 36: 1311–1318. Associação Brasileira de Divulgação Científica.

79. Tosi, A.J., Morales, J.C. & Melnick, D.J. 2000. Comparison of Y Chromosome and mtDNA Phylogenies Leads to Unique Inferences of Macaque Evolutionary History. Molecular Phylogenetics and Evolution 17: 133–144.

80. Tosi, A.J., Morales, J.C. & Melnick, D.J. 2003. Paternal, Maternal, and Biparental Molecular Markers Provide Unique Windows Onto the Evolutionary History of Macaque Monkeys. Evolution 57: 1419–1435.

81. Turkkahraman, D., Guran, T., Ivison, H., Griffin, A., Vijzelaar, R. & Krone, N. 2015. Identification of a Novel Large CYP17A1 Deletion by MLPA Analysis in a Family with Classic 17α-Hydroxylase Deficiency. SXD 9: 91–97. Karger Publishers.

82. de Waal, F.B.M. 2004. Peace Lessons from an Unlikely Source. PLOS Biology 2: e101. Public Library of Science.

83. Wang, S., Lawless, J. & Zheng, Z. 2020. Prenatal low-dose methyltestosterone, but not dihydrotestosterone, treatment induces penile formation in female mice and guinea pigs†. Biology of Reproduction 102: 1248–1260.

84. Witchel, S.F. 2018. DISORDERS OF SEX DEVELOPMENT. Best Pract Res Clin Obstet Gynaecol 48: 90– 102.

85. Xia, J., Liu, F., Wu, J., Xia, Y., Zhao, Z., Zhao, Y., et al. 2021. Clinical and Genetic Characteristics of 17 α-Hydroxylase/17, 20-Lyase Deficiency: c.985_987delTACinsAA Mutation of CYP17A1 Prevalent in the Chinese Han Population. Endocrine Practice 27: 137–145. Elsevier.

86. Xue, C., Raveendran, M., Harris, R.A., Fawcett, G.L., Liu, X., White, S., et al. 2016. The population genomics of rhesus macaques (Macaca mulatta) based on whole-genome sequences. Genome Res. 26: 1651–1662.

87. Yan, G., Zhang, G., Fang, X., Zhang, Y., Li, C., Ling, F., et al. 2011. Genome sequencing and comparison of two nonhuman primate animal models, the cynomolgus and Chinese rhesus macaques. Nature Biotechnology 29: 1019–1023.

88. Zhang, M., Sun, S., Liu, Y., Zhang, H., Jiao, Y., Wang, W., et al. 2015. New, recurrent, and prevalent mutations: Clinical and molecular characterization of 26 Chinese patients with 17alpha-hydroxylase/17,20-lyase deficiency. The Journal of Steroid Biochemistry and Molecular Biology 150: 11–16.

