## Supplementary Methods and Results for "Evolution of genes involved in the unusual genitals of the bear macaque, *Macaca arctoides*"

#### *Alignment, genome analysis, and variant calling*

For alignment, bwa-mem version 0.7.12 (Li & Durbin, 2009; Li, 2013) was used to separately align each read group and/or SRA file to the reference. After initial alignment to the reference, binary alignment map (bam) files for each sample were merged into a single file for each sample. Duplicate reads were then marked using Picard Tools version 1.79. Next, local re-alignment near insertion/deletion variants was completed using GATK's IndelRealigner (McKenna *et al.*, 2010). Primarily, we used GATK version 3.4-46 throughout, though we also used version 3.7 for VQSR and other subsequent steps. A summary of quality statistics for each sample throughout the genome analysis pipeline can be found in Table S1.

Following indel realignment, Base Quality Score Recalibration (BQSR) was performed using an iterative method due to the lack of a dbSNP for these samples. At this stage, samples from the same species were also merged into a single species-level bam file. First, an initial round of variant calling using GATK's Haplotype Caller was completed for each species and hard-filtered for high quality variants using vcftools (Danecek *et al.*, 2011) (Thresholds: minimum base quality: 40; minimum coverage: 15; maximum coverage: 160; minimum variant quality: 50). The base quality scores were then recalibrated, with each resulting recalibrated bam file run through HaplotypeCaller and hard filtered. Next, the new set of filtered variants were compared to the previous set of high quality variants using *vcf-compare* in vcftools. This iterative approach was repeated until fewer than 1% new variants were discovered, which resulted in two recalibration steps per sample. Visual inspection of the recalibration plots generated by BQSR revealed marginal improvement from the 1$^{st}$ and 2$^{nd}$ recalibration step. Average unique variants for step 1 were 1.98% (Table S1). The highest percentage for the 2$^{nd}$ recalibration step was 0.1% in *M. assamensis* samples (Table S1). Therefore, a less stringent 3% cutoff for new variants would have likely been sufficient for BQSR recalibration.

Following recalibration of base quality scores, SNP and indel variants were called together using GATK's Haplotype Caller for each sample. Following the last round of variant calling, we performed Variant Quality Score Recalibration (VQSR) of the quality values output by HaplotypeCaller. First, to generate a set of confident sites that are polymorphic, we generated two different training sets. We downloaded 389 whole genome sequences (WGS) of captive Indian Macaques from the Macaque Genotype and Phenotype Resource (ONPRC, n.d.) (mGAP, supported by NIH R24OD021324), some of which were previously described (Xue *et al.*, 2016). We also downloaded 81 WGS of wild-caught Chinese Macaques (Liu *et al.*, 2018) (Table S2).

We merged all files from mGAP and restricted only to biallelic SNPs. We similarly restricted the samples from Liu et al. (2018) to biallelic SNPs. The sites we obtained from Liu et al. (2018) form a training set of slightly less confident sites, while the mGAP sites will form a set of our most confident polymorphic sites. The mGAP database not only follows GATK best practices to call variants, but also exploits known pedigrees to check for Mendelian inconsistencies.

We ran the GATK VariantRecalibrator separately for each species VCF, using the following settings. We trained the recalibrator with mGAP sites with a Q12 prior (93.69%) and setting known to false, training to true, and truth to true. We trained the recalibrator with the Xue, et al. (2016) sites with a Q10 prior (90%) and setting known to false, training to true, and truth to false. We used the following annotations: QD, MQ, MQRankSum, ReadPosRankSum, FS, SOR, DP, and InbreedingCoeff. Finally, we used the GATK ApplyRecalibration module to filter all sites at the 99.9% tranche. This left us with final polymorphic site counts as summarized in Table 1. From each species-specific VQSR pass set, we used *vcf-compare* in vcftools to generate intersections of variants in all combinations across the five samples. We then used the R package UpSetR to generate an intersection plot (Figure 1C). Next, bedtools was used to get an intersection of the VQSR callsets for each species.

Because variants were called within each set of samples per species, there were non-passed variant genotypes for some species that did not pass VQSR. Therefore, we developed a separate filtering for variant and invariant sites which were eventually merged into a final callset. First, the raw gvcfs for each species from Haplotype Caller were jointly genotyped using GATK's joint genotype caller tool with the 'allSites' output option in reference confidence mode. Using vcftools, each species was subsetted and the VQSR passed sites were used to extract the variant callset for each species from this joint genotype output.

In order to compare invariant sites in the other samples for each variant position, potentially invariant genotypes were identified based on the intersected bedfile of VQSR pass sites. These sites were filtered from the jointly called set of genotypes using vcftools. The potentially invariant sites were filtered for variants (that did not pass VQSR filtering) and filtered based on depth (minimum: 5) and genotype quality (minimum: 20) using bcftools, which marks genotypes not matching criteria as missing data. Next, the variant and invariant callsets were concatenated and sorted using vcftools. Lastly, the separate species files were merged into a single filtered callset for all samples.

#### *Retrieving the outgroup base positions in papAnu4*

Finally, to get the corresponding base for the outgroup for each site, the masked baboon reference genome for papAnu4 was downloaded from UCSC. Both rheMac8 and papAnu4 were subsetted to chromosomes 1-20, chrX, and chrM for two-way alignment using lastz (--identity=75 --notransition --step=10 --gapped --chain --gfextend --format=maf+). The maf output was converted to a VCF-like output for baboon using a custom perl script, which included the alignment score, percent identity and orientation in the output. To filter overlapping alignments, the VCF-like file was sorted by position then by alignment score, keeping only the top scoring alignment for each position. Finally, all gap sites were removed.

#### *Comparison to genome analysis in original publications*

Compared to previous analyses using these samples, our genome analysis had a slightly lower coverage and higher duplication rate (Fan *et al.*, 2018), perhaps due to mapping to the masked reference genome, which reduced the possible callset size. However, this stringent masking likely resulted in an overall improvement in the quality of the sites we were able to call. When comparing the publicly available samples, we estimated slightly higher heterozygosity per sample compared to previous studies and slightly lower overall variation, indicating that our variant workflow is more stringent. We observed the lowest heterozygosity for *M. thibetana*, and the highest heterozygosity for *M. assamensis*, with *M. arctoides* having intermediate levels of heterozygosity.

#### *Complex speciation of the bear macaque*

Macaques are a diverse primate genus with several examples of purported complex speciation (Tosi *et al.*, 2000, 2003; Zinner *et al.*, 2011). Although mtDNA suggests that introgression with the *fascicularis* group should be driven by *M. mulatta* rather than *M. fascicularis* (Fan *et al.*, 2018; Roos *et al.*, 2019), an explicit test considering these two species separately found that the autosomal estimates of $f_{dM}$ were similar (Table S3). One possible explanation for this is that the sample used for *M. fascicularis* is Vietnamese in origin and therefore has extensive gene flow from *M. mulatta* (Yan *et al.*, 2011). It is worth noting that Fan *et al*. (2018) did a similar explicit test, but with *M. fascicularis* samples with little to no *M. mulatta* admixture and found no admixture signal. Similarly, our data found 1.6 times more shared heterozygous sites between *M. arctoides* and *M. mulatta* than between *M. arctoides* and *M. fascicularis* (Figure 2C). We also found 11.07% of genomic regions support a clustering of *M. arctoides* separate from the *fascicularis* and *sinica* species groups. Although several taxonomic surveys have suggested the placement of *M. arctoides* into its own species group (Fooden, 1980; Melnick & Kidd, 1985; Roos *et al.*, 2014), recent genome surveys suggest that sequencing of additional *sinica* species members will be useful in making this determination (Roos, 2018; Hettiarachchi, Nilmini *et al.*, 2019).

While genomic mosaicism is a key feature of species of hybrid origin, it is also a ubiquitous feature of admixture more generally, including when species initially emerge via bifurcating speciation. For example, in swordtail fish, there is genomic mosaicism that resembles hybrid speciation, although it is not consistent with other information from this species (Schumer *et al.*, 2014, 2016, 2018). Similarly, while a majority of the genome supports clustering of *M. arctoides* with *sinica* species, nearly 16% of the genome supports clustering of *M. arctoides* with *fascicularis* species. This deviation from a 50-50 split of ancestry from the two groups by itself does not necessarily contradict a hybrid origin, as it is not a requirement of hybrid speciation that each parental species contribute an equivalent genetic proportion to a species of hybrid origin  (e.g. Heliconius butterflies Jiggins *et al.*, 2008; but see Mavárez *et al.*, 2021). Here, this difference in parental contribution could be due to a combination of proposed recent admixture with *sinica* group taxa (Tosi *et al.*, 2003) and differences in ancestral populations sizes of the putative parental taxa. Subsequent gene flow with parental taxa poses an additional layer of complexity in this speciation scenario that makes the use of existing statistics that explicitly test for HHS (Hibbins & Hahn, 2019), not applicable due to a restrictive set of underlying assumptions. Still, this finding is also not uncommon in other taxa that do not have hybrid speciation. For example, in human-chimp-gorilla comparisons, ~30% of the human genome does not support the species tree (Scally *et al.*, 2012).

Because neither mosaicism or differences in parental contribution definitively support or reject HHS, nucleotide divergence was calculated based on the three possible topologies to investigate the evolution of the bear macaque further (Figure S5). These results revealed that the interspecies divergence between sister taxa in topo2 and topo3 regions were significantly different from each other (p=0), whereas hybrid speciation would predict similar divergence between these regions. Interestingly, rather than a deeper divergence between *M. arctoides* and the *sinica* group species, our results show the opposite. This higher divergence of topo3 regions than topo2 regions suggests that *M. arctoides* was initially more closely related to the *fascicularis* species group than to the *sinica* group. One explanation is that shortly after the split between *M. arctoides* and proto-f*ascicularis*, *M. arctoides* hybridized extensively with proto-*sinica* species, which resulted in nuclear swamping (Figure S1B). Another possibility is that perhaps *M. arctoides* diverged prior to the split between the *sinica* and *fascicularis* species groups, and subsequently had gene flow with the proto-*fascicularis* ancestor, followed by gene flow with the proto-*sinica* ancestor (Figure S1C). Both of these alternative scenarios explain the discordant mitochondrial and nuclear phylogenetic relationships (see Introduction), the higher proportion of regions supporting clustering with the *sinica* group (Figure 2D and Figure 3) as well as our divergence results (Figure S5).

### Supplementary Figures

**Supplementary Figure 1.** (A) Model based on literature (Tosi *et al.*, 2003; Jiang *et al.*, 2016; Fan *et al.*, 2018) depicts possible hybrid speciation between sinica and fascicularis species groups to form *M. arctoides*, with split time uncertainty relative to split of fascicularis group species, *M. mulatta* and *M. fascicularis* (Roos *et al.*, 2019). Approximate split times from (Jiang *et al.*, 2016). For this scenario, divergence in topo3 should be equal to that of topo2 (see Figure 5). Two alternative speciation scenarios for the bear macaque (B and C) based on divergence analysis results which are not consistent with Panel A (see Figure 5). Instead, our results suggest either that gene flow between *M. arctoides* and the sinica group members followed an older split with a proto-fascicularis ancestor (B). Alternatively, *M. arctoides* could have formed prior to the split between the two species groups and had subsequent gene flow first with the proto-fascicularis ancestor and then with the proto-sinica ancestor (C). Both of these alternative models contrast previous literature (A) and would predict a relative amount of divergence between sister taxa to be higher in topo3 than topo2.

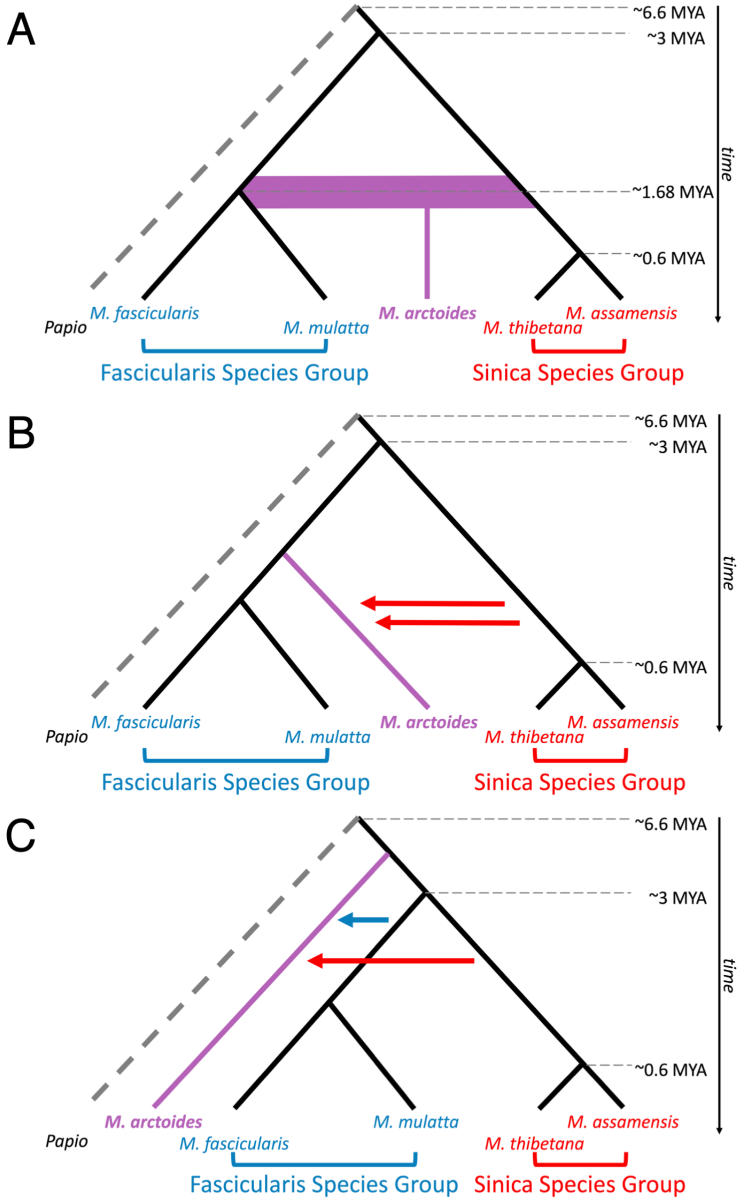

**Supplementary Figure 2. Introgression Analysis Methodology.** Below is a diagram of how the $f_{dM}$ statistic was calculated. First, the baboon reference genome was used to identify sites where *M. arctoides* had an alternative allele. Among these subsetted sites, an excess of derived shared alleles between *M. arctoides* and members of the sinica species group resulted in a negative $f_{dM}$ value. Alternatively, an excess of derived shared alleles between *M. arctoides* and members of the fascicularis species group resulted in a positive values $f_{dM}$ value. Values on either end of this distribution were targeted. In order to target sites that were unique in the *M. arctoides* lineage, the topology results were used. This is because a value in the middle of this distribution does not necessarily reflect sites that are unique to the *M. arctoides* lineage, only that there is not an excess in either direction.

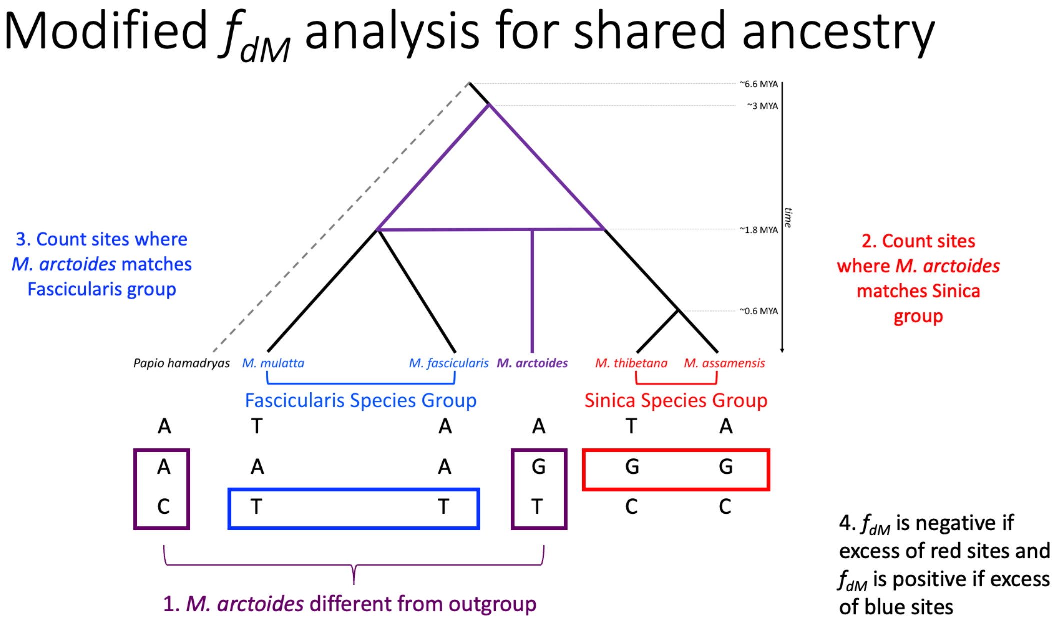

**Supplementary Figure 3.** Results from (a) 25kb, (b) 100kb, (c) 500kb, and (d) 1Mb sliding window analysis of $f_{dM}$. Regions where *M. arctoides* has sinica ancestry are displayed as negative values and regions where *M. arctoides* shares ancestry with fascicularis are displayed as positive values. Additionally, we excluded *M. arctoides* and made similar plots for each of the four other taxa at 50kb, none of which show the same level of mosaicism represented in *M. arctoides*.

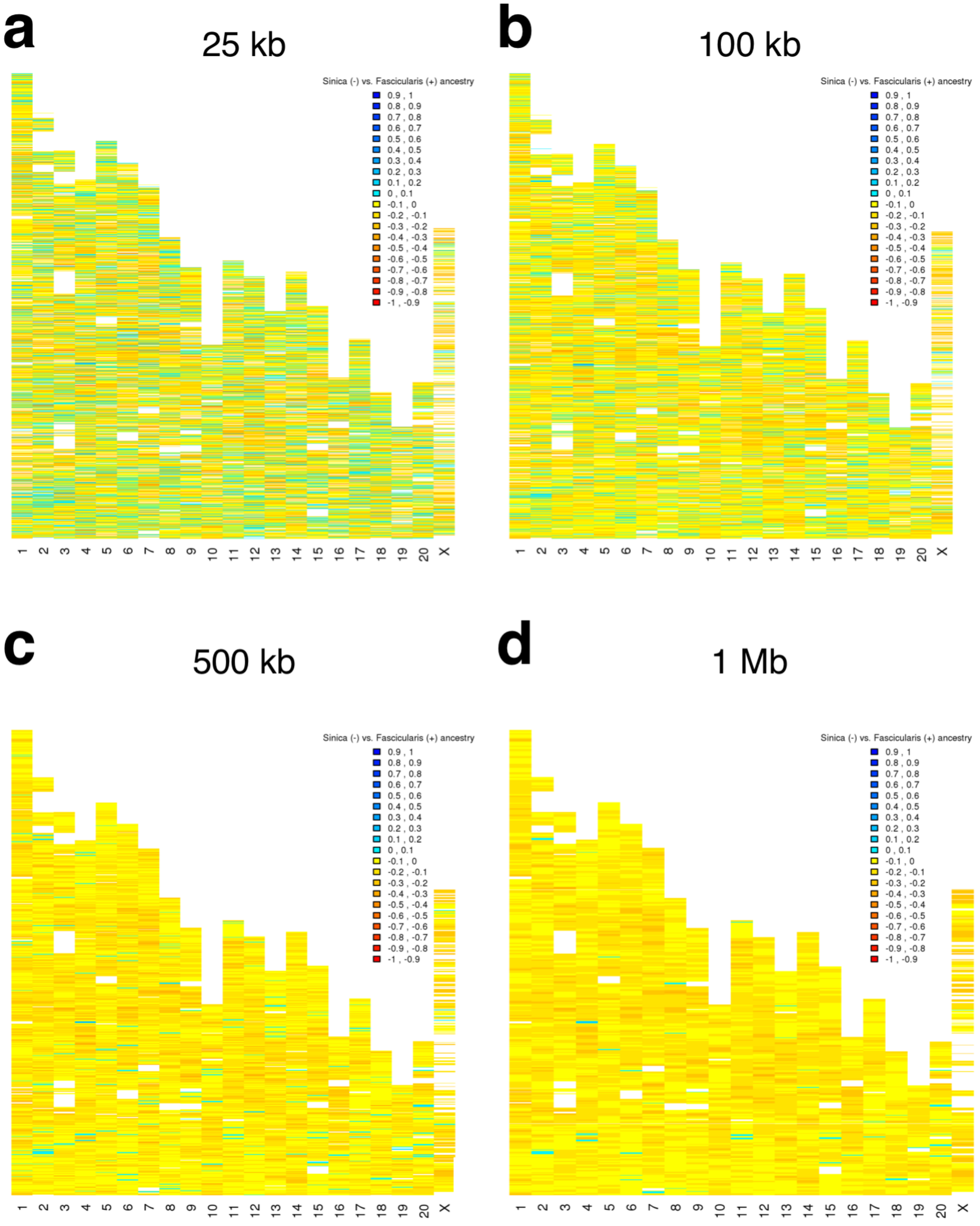

**Supplementary Figure 4.** Results from analysis of introgression (*fdM*) in 50kb sliding windows similar to Figure 3, except repeated using individual *M. arctoides* samples. Panel (a) shows only the Malaya sample, panel (b) shows only the SM1 sample, and panel (c) shows only the SM2 sample. As in Figure 3, regions where *M. arctoides* has most recent ancestry with the sinica group are displayed as negative values and regions where *M. arctoides* shares most recent common ancestry with the fascicularis group are displayed as positive values.

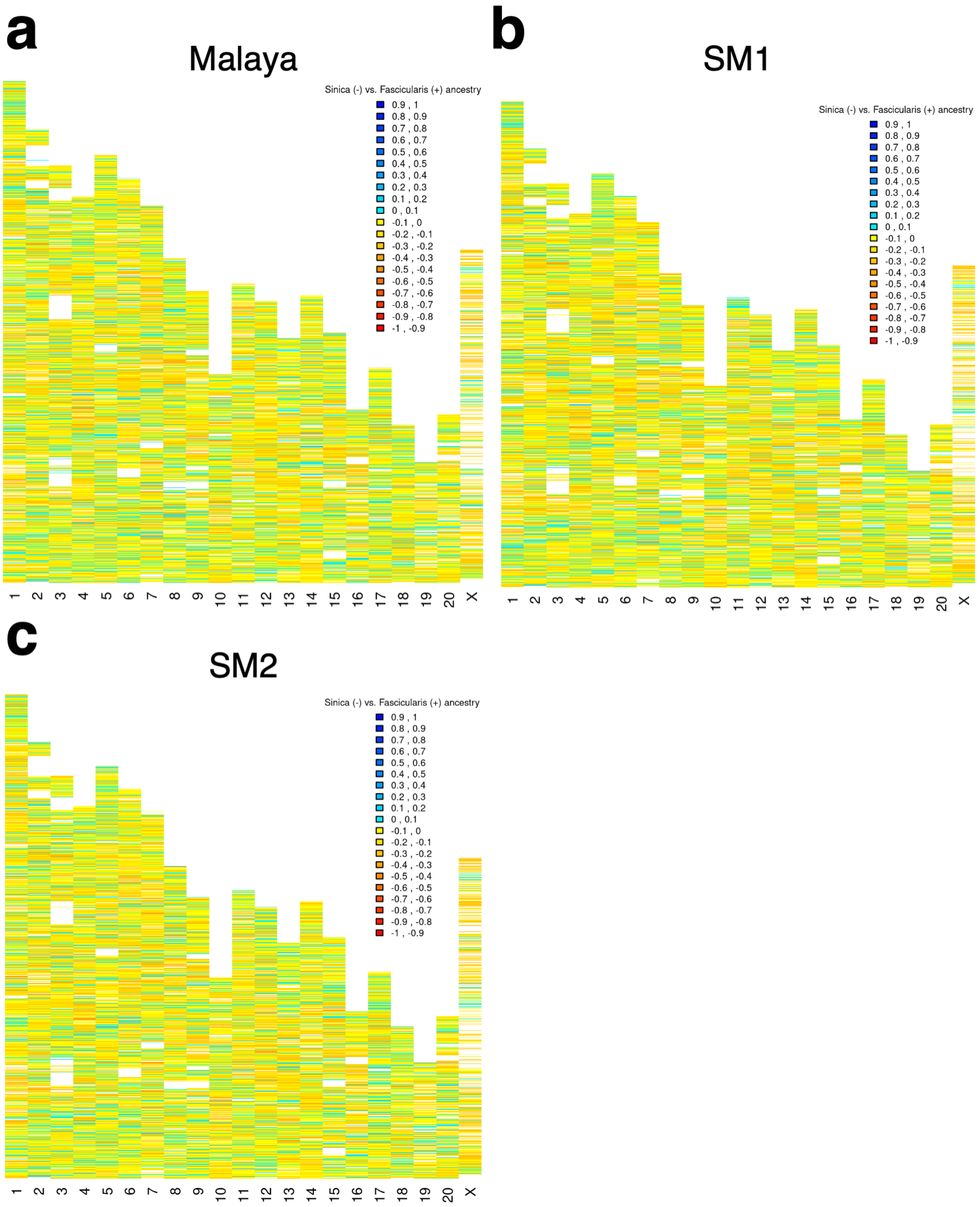

**Supplementary Figure 5. Nucleotide divergence casts doubt on hybrid speciation.** Nucleotide divergence, $D_{XY}$, was calculated between *M. arctoides* and its closest ancestor for regions of each of the three possible gene trees (see Figure 2D). These results reveal significant differences with topo1 having the deepest divergence consistent with ancestral polymorphism, and topo3 having the 2nd deepest, suggesting a more recent common ancestor for the *fascicularis* species group and the bear macaque than the *sinica* species group. Finally, the lowest divergence was in topo2 regions. Because the distribution of divergence is different between topo2 and topo3, this is not consistent with a hybrid speciation scenario (Figure S1A) where divergence should be the same. Instead, it suggests two alternative possible speciation scenarios shown in Figure S1B-C.

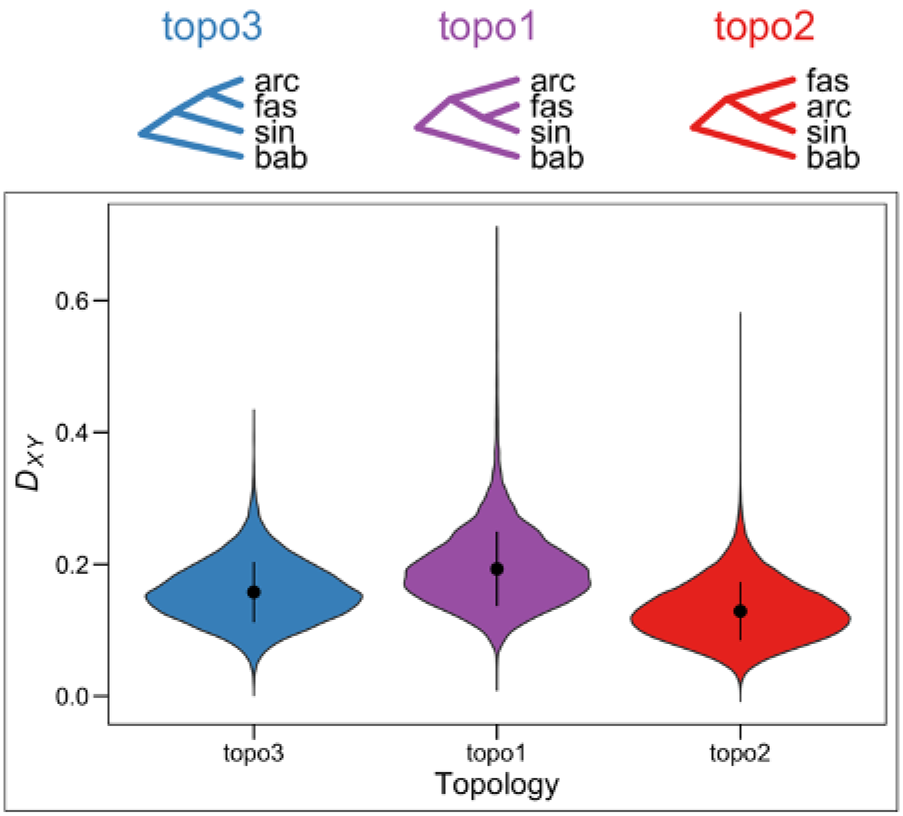

### Supplementary References


Danecek, P., Auton, A., Abecasis, G., Albers, C.A., Banks, E., DePristo, M.A., *et al.* 2011. The variant call format and VCFtools. *Bioinformatics* **27**: 2156–8.

Fan, Z., Zhou, A., Osada, N., Yu, J., Jiang, J., Li, P., *et al.* 2018. Ancient hybridization and admixture in macaques (genus Macaca) inferred from whole genome sequences. *Mol. Phylogenet. Evol.* **127**: 376–386.

Fooden, J. 1980. Classification and Distribution of Living Macaques (Macaca Lacepede, 1799). In: *The Macaques: Studies in Ecology, Behavior, and Evolution* (D. G. Lindburg, ed), pp. 1–9. Van Nostrand Reinhold Company, New York.

Hettiarachchi, Nilmini, Osada, Naoki, Nakaoka, Hirofumi, Hayakawa, Takashi, Inoue, Ituro, Saitou, Naruya, *et al.* 2019. Exome sequences of toque macaques (Macaca sinica) of Sri Lanka reveal many amino acid changes | bioRxiv. *bioRxiv*, doi: 10.1101/740761.

Hibbins, M.S. & Hahn, M.W. 2019. The Timing and Direction of Introgression Under the Multispecies Network Coalescent. *Genetics* **211**: 1059–1073.

Jiang, J., Yu, J., Li, J., Li, P., Fan, Z., Niu, L., *et al.* 2016. Mitochondrial Genome and Nuclear Markers Provide New Insight into the Evolutionary History of Macaques. *PLoS ONE* **11**.

Jiggins, C.D., Salazar, C., Linares, M. & Mavarez, J. 2008. Hybrid trait speciation and Heliconius butterflies. *Philos. Trans. R. Soc. B Biol. Sci.* **363**: 3047–3054.

Li, H. 2013. Aligning sequence reads, clone sequences and assembly contigs with BWA-MEM. *arXiv* **1303**.

Li, H. & Durbin, R. 2009. Fast and accurate short read alignment with Burrows-Wheeler transform. *Bioinformatics* **25**: 1754–1760.

Liu, Z., Tan, X., Orozco-terWengel, P., Zhou, X., Zhang, L., Tian, S., *et al.* 2018. Population genomics of wild Chinese rhesus macaques reveals a dynamic demographic history and local adaptation, with implications for biomedical research. *GigaScience* **7**.

Mavárez, J., Salazar, C.A., Bermingham, E., Salcedo, C., Jiggins, C.D. & Linares, M. 2021. Author Correction: Speciation by hybridization in Heliconius butterflies. *Nature* **592**: E4–E5.

McKenna, A., Hanna, M., Banks, E., Sivachenko, A., Cibulskis, K., Kernytsky, A., *et al.* 2010. The Genome Analysis Toolkit: a MapReduce framework for analyzing next-generation DNA sequencing data. *Genome Res* **20**: 1297–303.

Melnick, D.J. & Kidd, K.K. 1985. Genetic and evolutionary relationships among Asian Macaques. *Int. J. Primatol.* **6**: 123–160.

ONPRC. n.d. mGAP: The Macaque Genotype And Phenotype Resource.

Roos, C. 2018. Complete mitochondrial genome of a Toque Macaque (Macaca sinica). *Mitochondrial DNA Part B* **3**: 182–183. Taylor & Francis.

Roos, C., Boonratana, R., Supriatna, J., Fellowes, J.R., Groves, C.P., Nash, S.D., *et al.* 2014. An Updated taxonomy and Conservation Status Review of Asian Primates. *Asian Primates J. Vol. 4 1 2-38*. IUCN/SSC Primate Specialist Group.

Roos, C., Kothe, M., Alba, D.M., Delson, E. & Zinner, D. 2019. The radiation of macaques out of Africa: Evidence from mitogenome divergence times and the fossil record. *J. Hum. Evol.* **133**: 114–132.

Scally, A., Dutheil, J.Y., Hillier, L.W., Jordan, G.E., Goodhead, I., Herrero, J., *et al.* 2012. Insights into hominid evolution from the gorilla genome sequence. *Nature* **483**: 169–175. Nature Publishing Group.

Schumer, M., Cui, R., Powell, D.L., Rosenthal, G.G. & Andolfatto, P. 2016. Ancient hybridization and genomic stabilization in a swordtail fish. *Mol. Ecol.* **25**: 2661–2679.

Schumer, M., Rosenthal, G.G. & Andolfatto, P. 2014. How Common Is Homoploid Hybrid Speciation?

*Evolution* **68**: 1553–1560.
Schumer, M., Rosenthal, G.G. & Andolfatto, P. 2018. What do we mean when we talk about hybrid speciation? *Heredity* **120**: 379.
Tosi, A.J., Morales, J.C. & Melnick, D.J. 2000. Comparison of Y Chromosome and mtDNA Phylogenies Leads to Unique Inferences of Macaque Evolutionary History. *Mol. Phylogenet. Evol.* **17**: 133–144.
Tosi, A.J., Morales, J.C. & Melnick, D.J. 2003. Paternal, Maternal, and Biparental Molecular Markers Provide Unique Windows Onto the Evolutionary History of Macaque Monkeys. *Evolution* **57**: 1419–1435.
Xue, C., Raveendran, M., Harris, R.A., Fawcett, G.L., Liu, X., White, S., *et al.* 2016. The population genomics of rhesus macaques (Macaca mulatta) based on whole-genome sequences. *Genome Res.* **26**: 1651–1662.
Yan, G., Zhang, G., Fang, X., Zhang, Y., Li, C., Ling, F., *et al.* 2011. Genome sequencing and comparison of two nonhuman primate animal models, the cynomolgus and Chinese rhesus macaques. *Nat. Biotechnol.* **29**: 1019–1023.
Zinner, D., Arnold, M.L. & Roos, C. 2011. The strange blood: Natural hybridization in primates. *Evol. Anthropol. Issues News Rev.* **20**: 96–103.
